# Glycosylation of bioactive C_13_-apocarotenols in *Nicotiana benthamiana* and *Mentha × piperita*

**DOI:** 10.1101/2020.07.29.225110

**Authors:** Guangxin Sun, Natalia Putkaradze, Sina Bohnacker, Rafal Jonczyk, Tarik Fida, Thomas Hoffmann, Rita Bernhardt, Katja Härtl, Wilfried Schwab

**Affiliations:** Biotechnology of Natural Products, Technische Universität München, Liesel-Beckmann-Str. 1, 85354 Freising, Germany; Institut für Biochemie, Universität des Saarlandes, Campus B2 2, D-66123 Saarbrücken, Germany

**Keywords:** glycosyltransferase, apocarotenoid, allelochemical, *Mentha × piperita*, *Nicotiana benthamiana*, cytochrome P450

## Abstract

C_13_-apocarotenoids (norisoprenoids) are carotenoid-derived oxidation products, which perform important physiological functions in plants. Although their biosynthetic pathways have been extensively studied, their metabolism including glycosylation remains elusive. Candidate uridine-diphosphate glycosyltransferase genes (*UGTs*) were selected for their high transcript abundance in comparison with other *UGTs* in vegetative tissues of *Nicotiana benthamiana* and *Mentha × piperita*, as these tissues are rich sources of apocarotenoid glucosides. Hydroxylated C_13_-apocarotenol substrates were produced by P450-catalyzed biotransformation and microbial/plant enzyme systems were established for the synthesis of glycosides. Natural substrates were identified by physiological aglycone libraries prepared from isolated plant glycosides. In total, we identified six UGTs that catalyze the unprecedented glucosylation of C_13_-apocarotenols, where glucose is bound either to the cyclohexene ring or butane side chain. MpUGT86C10 is a superior novel enzyme that catalyzes the glucosylation of allelopathic 3-hydroxy-*α*-damascone, 3-oxo-*α*-ionol, 3-oxo-7,8-dihydro-*α*-ionol (Blumenol C) and 3-hydroxy-7,8-dihydro-*β*-ionol, while a germination test demonstrated the higher phytotoxic potential of a norisoprenoid glucoside in comparison to its aglycone. Glycosylation of C_13_-apocarotenoids has several functions in plants, including increased allelopathic activity of the aglycone, facilitating exudation by roots and allowing symbiosis with arbuscular mycorrhizal fungi. The results enable in-depth analyses of the roles of glycosylated norisoprenoid allelochemicals, the physiological functions of apocarotenoids during arbuscular mycorrhizal colonization and the associated maintenance of carotenoid homeostasis.

**One-sentence summary:** We identified six transferases in *Nicotiana benthamiana* and *Mentha x piperita*, two rich sources of glycosylated apocarotenoids that catalyze the unprecedented glycosylation of a range of hydroxylated α- and β-ionone/ionol derivatives and were able to modify bioactivity by glucosylation.

## Introduction

Plants synthesize a number of C_40_ lipid-soluble colorful carotenoids and oxygen-bearing xanthophylls from C_5_ isopentenyl building blocks, which are essential for photosynthesis and –protection (Giuliano 2014; Tian 2015). They occur in all photosynthetic organisms (higher plants, algae, and cyanobacteria) as well as some non-photosynthetic microbes (fungi and bacteria) (Cazzonelli and Pogson 2010; Walter and Strack 2011; Zhang 2018). When produced in petals and other parts of flowers carotenoids/xanthophylls act as visual signals to attract pollinators, while they decoy seed-dispersing animals when accumulated in fruits. Therefore, C_40_-isoprenoids are also essential for plant reproduction (Wurtzel 2019). Chloroplast-associated carotenoids stabilize membranes, and are required to form prolamellar bodies (Park et al. 2002).

Besides, carotenoids/xanthophylls are precursors of apocarotenoids, which are formed by carotenoid cleavage oxygenases (CCOs) and have important functions in plant development, growth, architecture, and plant-environment interactions such as the attraction of pollinators and the defense against pathogens and herbivores (Hou et al. 2016; Nisar et al. 2015; Ohmiya et al. 2006; Ohmiya 2009; Tian 2015; Walter et al. 2010). More precisely, bioactive apocarotenoids act as hormones, signaling compounds, allelopathic substances, chromophores, scent/aroma constituents, repellents, chemoattractants, growth stimulators and inhibitors. They comprise the C_20_-vitamin A derivatives (retinal, retinol, and retinoic acid), the C_20_-saffron pigment crocetin, the C_15_-phytohormone abscisic acid (ABA), strigolactones, volatile (C_9_ and C_13_) and non-volatile degradation products (Dickinson et al. 2019; Finkelstein 2013; Hou et al. 2016; Walter et al. 2010; Zhang 2018). There are at least two types of CCOs, the 9-*cis* epoxycarotenoid dioxygenases (NCEDs) that catalyze the first step in ABA biosynthesis, and carotenoid cleavage dioxygenases (CCDs) that specifically oxidize carotenoids at different double bonds leading to metabolites of different sizes (Huang et al. 2009; Nisar et al. 2015). In plants, apocarotenoids accumulate particularly in certain plastids (etioplasts, leucoplasts and chromoplasts) and tissues such as flowers, leaves and roots (Lohse et al. 2005; Strack and Fester 2006).

ABA was the first apocarotenoid to be discovered in plants, and the enzymes for biosynthesis and degradation of ABA are identified and quite well characterized (Finkelstein 2013). However, in addition to ABA plants biosynthesize many more apocarotenoids but their mechanisms of action, biochemical modifications, associated enzymes, regulation, and transporters remain elusive (Finkelstein 2013; Hou et al. 2016). The total number of apocarotenoids and associated bioactivities is largely unknown but they help fine-tune carotenogenesis, plant development and environmental responses (Hou et al. 2016; Lätari et al. 2015). There are still numerous unknown apocarotenoids that function as signaling compounds to control plant architecture, since blocking carotenoid biosynthesis or eliminating CCDs led to architectural anomalies (Dickinson et al. 2019; Hou et al. 2016). Furthermore, changes in apocarotenoid accumulation in response to developmental and environmental cues demonstrate that some degradation products have regulatory roles in plants (Avendaño-Vázquez et al. 2014). These recent discoveries of apocarotenoid bioactivities indicate an untapped potential for plant modification to meet the needs of agriculture and industry.

C_13_-apocarotenoids, also known as norisoprenoids, are 13-carbon butene cyclohexene degradation products formed by the cleavage of carotenoids (Supplemental Figure S1) (Winterhalter and Rouseff 2002). Many of them are volatile and contribute to the flavor and aroma of flowers and fruits. These volatiles are highly valued by industry due to their low odor threshold values, and characteristic aroma notes (e.g. *α*-ionone, *β*-ionone, *α*-damascone, and *β*-damascone (Cataldo et al. 2016; Rodríguez-Bustamante and Sánchez 2007; Walter and Strack 2011). The biological functions of norisoprenoids go beyond the frequently discussed attraction of seed dispersers and pollinators as *β*-ionone and some other apocarotenoids such as 3-oxo-7,8-dihydro-*α*-ionone/ionol show also antimicrobial and antifungal activity (Park et al. 2004; Walter and Strack 2011). Tobacco plants, infected by blue mold accumulated *β*-ionone levels 50–600-fold higher in non-infected stem tissues adjacent to necrotic lesions (Salt et al. 1986), and many norisoprenoids with hydroxy- or oxo-functions at C3 position act as plant growth inhibitors and allelochemicals (D’Abrosca et al. 2004; Dietz and Winterhalter 1996; Kato-Noguchi et al. 2010; Macías et al. 2008). Blumenol (3-oxo-*α*-ionol and its glycosides) accumulates upon arbuscular mycorrhiza (AM) colonization and is probably responsible for systemic suppression of additional AM colonization (Hou et al. 2016; Wang et al. 2018). Bioactive apocarotenoids often undergo enzymatic transformations (Mathieu et al. 2009) such as oxidation, reduction, and glycosylation, which modify their biological activities. Therefore, norisoprenoids occur predominantly in bound forms, i.e., glycosylated (Cai et al. 2014; Çalış et al. 2002; Ito et al. 2000; Ito et al. 2001; Kodama et al. 1981; Neugebauer et al. 1995; Pabst et al. 1992b; Schwab and Schreier 1990; Tommasi et al. 1996; Wirth et al. 2001). Interestingly, C_13_-apocarotenoids occur almost exclusively as β-D-glucosides, but the second bound sugar can be different (Pabst et al. 1992a). Glycosylation of plant metabolites enhances their stability and water solubility, facilitates their storage and accumulation, reduces the toxicity of potential toxic agents, and is a key mechanism in the metabolic homeostasis of plant cells (Bowles et al. 2005).

In plants, uridine-diphosphate sugar depending glycosyltransferases (UGTs) catalyze the production of small molecule glycosides by transferring a carbohydrate from an activated monosaccharide donor, usually UDP-glucose, to an alcohol, acid, amine, or thiol (Song et al. 2018; Wang and Hou 2009). Genomes of higher plants have more than 100 UGTs (Caputi et al. 2012). Plant UGTs acting on small molecules are mostly members of the carbohydrate active enzyme (CAZy; http://www.cazy.org) GT family 1 (UDP sugar-dependent UGTs) consisting of GT-B fold enzymes (Song et al. 2018). Family 1 UGTs are majorly involved in glycosylation of terpenoids, alkaloids, cyanohydrins, glucosinolates, flavonoids, and phenylpropanoids (Asada et al. 2013; Augustin et al. 2012; Bönisch et al. 2014; Bowles et al. 2005). To date, only few UGTs are known to glycosylate apocarotenoids.

Crocin, an apocarotenoid glycosyl ester is produced by the sequential action of UGT75L6 and UGT94E5 in *Gardenia jasminoides* (Nagatoshi et al. 2012). UGT75L6 glucosylates the carboxyl group of crocetin yielding crocetin glucosyl esters, while UGT94E5 transfers glucose to the 6’ hydroxyl group of the glucose moiety of crocetin glucosyl esters. UGTCs2 from *Crocus sativus* has glucosylation activity against crocetin, crocetin *β*-D-glucosyl ester and crocetin *β*-D-gentibiosyl ester (Moraga et al. 2004) leading to highly glucosylated crocins (Ahrazem et al. 2015). Only recently, it was shown that UGT709G1 also from *C. sativus* catalyzes the glucosylation of 3-hydroxy-*β*-cyclocitral, making it suited for the biosynthesis of picrocrocin, the precursor of safranal (Diretto et al. 2019). However, UGTs reacting with C_13_-apocarotenoids are unknown.

*Nicotiana* spp are one of the richest sources of carotenoid degradation products, with nearly 100 components identified (Bolt et al. 1983; Ito et al. 2000; Takagi et al. 1980; Wahlberg and Enzell 1987). Although *N. benthamiana* is an indispensable research model that can be genetically modified efficiently (Goodin et al. 2008), little is known about the production of apocarotenoids in this plant system and in particular about UGTs transforming small molecules (Jassbi et al. 2017; Sun et al. 2019). Several draft assemblies of its genome have been generated and a recent database presented 42,855 putative genes (Kourelis et al. 2018) containing 174 UGT genes (http://supfam.org/SUPERFAMILY). Recently, we isolated ten *UGT* genes from *N. benthamiana* and characterized their encoded proteins (Sun et al. 2019). They showed promiscuity towards a number of plant metabolites including phenolics and terpenoids and were further investigated in this work for glucosylation of apocarotenoids.

The genus of mint (*Mentha*, Lamiaceae) includes approximately 25–30 species that are widely used as medicinal and aromatic herbs. Peppermint (*M. × piperita*) is a sterile (hexaploid) hybrid created from watermint (*M. aquatica*) and spearmint (*M. spicata* (Ahkami et al. 2015). An important product of the members of the mint genus is the essential oil, whose valuable ingredients are menthol and menthone (Croteau et al. 2005). While the biosynthesis of (−)-menthol and its isomers has been thoroughly analyzed (Croteau et al. 2005) and UGT products, such as flavone glycosides (Erenler et al. 2018) and menthol glycoside (Sgorbini et al. 2015) have been described in *Mentha* species, UGTs have not yet been characterized. Norisoprenoids have not yet been detected in *Mentha* species. A draft genome sequence of *M. longifolia*, a diploid species ancestral to cultivated peppermint and spearmint, is available (Vining et al. 2017).

Here, we describe the isolation, identification and characterization of unprecedented C_13_-apocarotenol UGTs from *N. benthamiana* and *M. × piperita*. The plant species were selected as potential sources for UGTs because tobacco is a known producer of allelopathic norisoprenoids (Bolt et al. 1983; D’Abrosca et al. 2004; Ito et al. 2000; Jassbi et al. 2017; Kodama et al. 1981; Kodama et al. 1984; Mushtaq and Siddiqui 2018). Mint was chosen as second source because related glycosides have been isolated from mint but no UGT has been characterized in this plant. Recombinant proteins were screened with the commercially available model C_13_-apocarotenols *α*- and *β*-ionol, and candidate UGTs were then used to glucosylate hydroxylated norisoprenoids (Supplemental Figure S1) produced by P450-mediated biotransformation. Aglycone libraries identified the natural substrates of C_13_-apocarotenol UGTs, while agroinfiltration and *β*-ionol application demonstrated the importance of substrate availability. Germination tests were performed to compare the phytotoxic effect of a model norisoprenoid glucoside and its respective aglycone. This knowledge of norisoprenoid UGTs can contribute significantly to the identification of unknown apocarotenoid signal compounds in plants and the function of their glucosides during arbuscular mycorrhizal fungi colonization of *Nicotiana* roots (Wang et al. 2018).

## Results

### Selection of candidate UGTs and protein expression

The study was undertaken to identify the first plant UGTs that glucosylate C_13_-apocarotenoids. UGTs from two plant species (N. *benthamiana* and *M. × piperita*) were characterized. Since C_13_-apocarotenoid glycosides accumulate in vegetative tissues (Lätari et al. 2015) and the tobacco plant is a rich source of theses metabolites, we selected ten *UGT* genes (*NbUGT71AJ1, NbUGT72AX1, NbUGT72AY1, NbUGT72B34, NbUGT72B35, NbUGT73A24, NbUGT73A25, NbUGT85A73, NbUGT85A74*, and *NbUGT709Q1*), which were recently isolated from *N. benthamiana* leaves due to their high transcript levels in vegetative tissues in comparison with other *UGTs* (Sun et al. 2019). In addition, three full-length *UGT* gene sequences were found in a *M. × piperita* transcriptome database of the Mint Genomics Resource at the Washington State University (http://langelabtools.wsu.edu/mgr/home) (Ahkami et al. 2015). They were successfully cloned from *M. × piperita* and designated *MpUGT86C10, MpUGT708M1*, and *MpUGT709C6.* The UGT identities were assigned by the UGT Nomenclature Committee (https://prime.vetmed.wsu.edu/resources/udp-glucuronsyltransferase-homepage). The mRNA used to generate the transcriptome database was isolated from glandular trichomes of *M. x piperita* at two leaf developmental stages. Leaves of 0.5 to 1.5 cm length (top two pairs from the top) were harvested as ‘immature leaves’ (Ahkami et al. 2015). The 5th leaf pair from the top constituted ‘mature leaves’. During nucleic acid extraction extra alleles, *MpUGT708M2* and *MpUGT709C7/8* were obtained. Gene expression analysis of the 13 putative *UGTs* (ten from tobacco and three from mint) confirmed their significant transcript levels in vegetative tissues except for *NbUGT71AJ1, NbUGT709Q1*, and *NbUGT85A74* (Supplemental Figure S2). Since C_13_-apocarotenoid glycosides have so far been isolated mainly from plant leaves, these genes were good candidates to encode for C_13_-apocarotenoid UGTs. For biochemical characterization of the encoded proteins, the *UGT* genes were amplified from leaf cDNA of *M. × piperita* and *N. benthamiana*, and cloned into the pGEX-4T-1 expression vector containing an N-terminal glutathione S-transferase (GST-fusion) tag. The fusion proteins were successfully produced in *E. coli* BL21 (DE3) pLysS, affinity purified and verified by SDS-PAGE (Supplemental Figure S3). Clear GST-UGT protein bands were visible at around 80 kDa in all elution fractions, in addition to a band of about 27 kDa for the free GST protein. Western blot analyses using anti-GST-antibody confirmed the identity of GST-UGT fusion proteins.

### Substrate screening with model apocarotenoids and product identification

The recombinant UGTs from *N. benthamiana* and *M. × piperita*, which were produced in *E. coli*, were subjected to *in vitro* substrate screening. The proteins were incubated with UDP-glucose as sugar donor and the commercially available model C_13_-apocarotenols *α*- and *β*-ionol, as well as 3-oxo-*α*-ionol (kindly provided by Y. Gunata, Montpellier, France) as acceptors. Only UDP-glucose was used as acceptor substrate for the screening because apocarotenoids are bound almost exclusively to β-D-glucose and other acceptors are rarely available. Products were analyzed by LC-MS (Figure 1). Of the 16 UGTs tested, six and four were able to glucosylate *α/β*-ionol and 3-oxo-*α*-ionol, respectively. The glucosides of *α*- and *β*-ionol were detected in the ion traces *m/z* 401 [M+HCOO]^-^ and 391 [M+Cl]^-^ (Supplemental Table S1). Similarly, 3-oxo-*α*-ionyl glucoside was found at *m/z* 415 [M+HCOO]^-^ and 405 [M+Cl]^-^. All substrates were also incubated with empty vector (control) protein extracts (Figure 1). No glucoside was produced in these control samples. The peak shapes imply the formation of diastereomeric mixtures, in the case of *α*-ionyl- and 3-oxo-*α*-ionyl glucoside. To confirm the identity of diastereomeric *α*-ionyl *β*-D-glucopyranoside, recombinant *E. coli* cells expressing MpUGT86C10 were used as whole-cell biocatalyst for the production of the C_13_-apocarotenyl glucoside according to (Effenberger et al. 2019). The NMR data confirmed the structure of the product (Supplemental Figure S4) and were in accordance with those of (Zeng et al. 2014). The splitting of the signal for H9/C9 (4.33/72.81 ppm and 4.24/75.45 ppm) in HSQC proved the presence of a racemate and the H1’/C1’ signal (4.15/100.56) showed the typical chemical shift for *β*-D-glucosides. The protein sequence analysis of the 16 candidate proteins revealed that the six UGTs that catalyze the glucosylation of C_13_-apocarotenoids belong to different UGT families (72, 73, 85, 86, and 709) and contain the characteristic features of functional UGTs (Supplemental Figures S5 and S6). MpUGT86C10 stands out because it is separated from the other sequences. Pairwise amino acid sequence analysis of norisoprenoid UGTs showed only identities of 22.1 to 36.1%, except for NbUGT73A24 und NbUGT73A25, which exhibited 94.5% (Supplemental Table S2). The proteins that glucosylate C_13_-apocarotenols are 476 to 485 amino acids long and contain the characteristic motifs of a functional UGT including the catalytically active His (position 20 in MpUGT86C10) and Asp (position 130), the plant secondary product glycosyltransferase (PSPG) box (position 350– 393), and the GSS motif (position 456-458; Supplemental Figure S6) (Sun et al. 2019). Only the six UGTs that glucosylated C_13_-apocarotenoids were further considered (Table 1).

**Table 1.** C_13_-Apocarotenol screening of UGTs by LC-MS. *In vitro* reaction products were analyzed by LC-MS. The color code shows increasing reactivity from white to green color (maximum activity in dark green corresponding to 100%; 100% indicates the enzyme with the highest activity towards the specific substrate). The chemical structure illustrates the derived preferred glycosylation sites of the UGTs.

| substrates | NbUGT72AY1 | NbUGT73A24 | NbUGT73A25 | NbUGT85A73 | MpUGT86C10 | MpUGT709C6 |
| --- | --- | --- | --- | --- | --- | --- |
| α-Ionol | 77 | 77 | 100 | 21 | 66 | 7 |
| β-Ionol | 51 | 86 | 100 | 26 | 57 | 23 |
| 3-Hydroxy-α-ionol | 73 | 55 | 100 | 7 | 96 | 2 |
| 4-Hydroxy-β-ionol | 27 | 90 | 100 | 2 | 81 | 9 |
| 3-Hydroxy-α-ionone | 1 | 2 | 4 | 1 | 100 | 0 |
| 4-Hydroxy-β-ionone | 6 | 12 | 67 | 2 | 100 | 4 |
| 3-Hydroxy-α-damascone | 4 | 20 | 50 | 29 | 100 | 0 |
| 4-Hydroxy-β-damascone | 0 | 0 | 9 | 15 | 100 | 43 |
| 3-Oxo-α-ionol | 100 | 37 | 25 | 0 | 45 | 0 |

NbUGT73A25 NbUGT73A24 NbUGT72AY1 MpUGT86C10
preferred glucosylations sites of UGTs
MpUGT86C10 NbUGT73A25

**Fig. 1.**
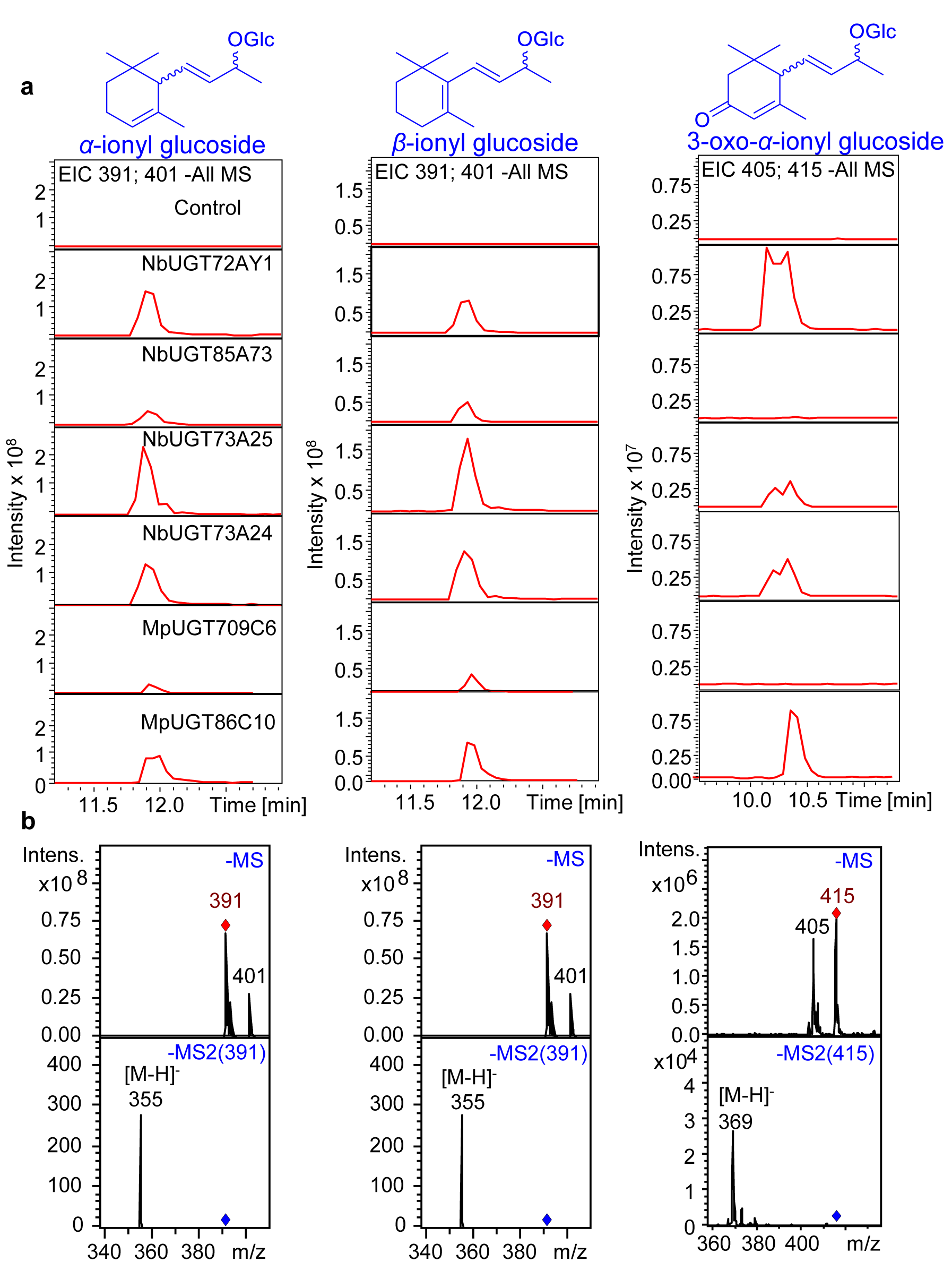
Identification of C_13_-apocarotenyl glucosides formed by UGTs from *N. benthamia* and *M. × piperita*. (**a**) Combined ion traces (extracted ion chromatogram EIC) *m/z* 391 [M+Cl]^-^ and *m/z* 401 [M+HCOO]^-^ (ionyl glucosides) as well as *m/z* 405 [M+Cl]^-^ and *m/z* 415 [M+HCOO]^-^ (3-oxo-α-ionyl glucoside). (**b**) mass spectra (-MS) and product ion spectra (-MS2). Diagnostic ions are explained in Supplemental Table S1. Glc glucopyranose.

### Kinetic analysis of recombinant proteins

First, the reaction conditions were optimized for each enzyme in terms of protein amount, incubation time, incubation temperature, and pH value (Supplemental Table S3). At least eight different concentrations (10-1,200 µM) of *α*- and *β*-ionol were used and the released UDP amounts were quantified to calculate the enzyme activities. For NbUGT72AY1, NbUGT73A24, NbUGT73A25, MpUGT86C10, and MpUGT709C6, the *K*_M_, *v*_max_, *k*_cat_ and *k*_cat_/*K*_M_ values were determined (Table 2). NbUGT85A73 showed only low activity and kinetic values were not measured. The *K*_M_ values ranged from 17.4 to 106.0 μM for *α*-ionol, and from 6.0 to 131.4 μM for *β*-ionol. Similarly, *k*_cat_ values ranged from 0.002 to 0.086 sec^-1^ for *α*-ionol, and 0.007 to 0.208 sec^-1^ for *β*-ionol. NbUGT73A25 and MpUGT86C10 showed similar and the highest enzyme specificity constants of 811 and 786 M^-1^ sec^-1^ for *α*-ionol, respectively and 1580 and 1575 M^-1^ sec^-1^ for *β*-ionol, respectively.

**Table 2.** Michaelis-Menten kinetic parameters of NbUGT72AY1, NbUGT73A24, NbUGT73A25, GT86C10, and MpUGT709C6. Kinetics were determined fora-ionol and /3-ionol using the UDP Glo™ assay.

| | Substrate | $K_M$ [μM] | $v_{max}$ [nmol min<sup>-1</sup> mg<sup>-1</sup>] | $k_{cat}$ [sec<sup>-1</sup>] | $k_{cat}/K_M$ [M<sup>-1</sup> sec<sup>-1</sup>] |
| --- | --- | --- | --- | --- | --- |
| NbUGT72AY1 | α-ionol | 17.4 ± 7.9 | 9.8 ± 0.7 | 0.013 | 749 |
| | β-ionol | 13.1 ± 6.8 | 5.0 ± 1.1 | 0.007 | 507 |
| NbUGT73A24 | α-ionol | 32.81 ± 8.8 | 18.5 ± 2.5 | 0.025 | 750 |
| | β-ionol | 34.01 ± 3.9 | 18.4 ± 0.2 | 0.024 | 718 |
| NbUGT73A25 | α-ionol | 106.0 ± 22.2 | 64.4 ± 4.5 | 0.086 | 811 |
| | β-ionol | 131.4 ± 18.3 | 155.6 ± 7.1 | 0.208 | 1580 |
| MpUGT86C10 | α-ionol | 23.8 ± 3.7 | 14.0 ± 0.8 | 0.019 | 786 |
| | β-ionol | 6.0 ± 2.1 | 7.0 ± 1.4 | 0.009 | 1575 |
| MpUGT709C6 | α-ionol | 27.2 ± 0.4 | 1.3 ± 0.1 | 0.002 | 65 |
| | β-ionol | 28.7 ± 7.5 | 7.2 ± 1.3 | 0.010 | 336 |

### Production of hydroxylated C_13_-apocarotenols and their conversion by UGTs

It has recently been shown that bacterial cytochrome P450s, among them the hydroxylase CYP109E1 from *Bacillus megaterium*, can oxidize various apocarotenoids (Khatri et al. 2010; Litzenburger and Bernhardt 2016) including ionones (Putkaradze et al. 2017). Therefore, *B. megaterium* expressing CYP109E1 was used in this study as whole-cell biocatalyst to produce hydroxylated C_13_-apocarotenoids from *α*-, and *β*-ionol, *α*- and *β*-ionone, and *α*-, and *β*-damascone to test them as potential substrates for the candidate UGTs as these C_13_-apocarotenols are not commercially available. The main products of the biotransformation using CYP109E1 were 3-hydroxy-*α*-ionol, 4-hydroxy-*β*-ionol, 3-hydroxy-*α*-ionone, 4-hydroxy-*β*-ionone, 3-hydroxy-*α*-damascone, and 4-hydroxy-*β*-damascone (Supplemental Figures S7-S12). Their identities were confirmed by GC-MS analysis and comparison with reference data (Schoch et al. 1991). The crude mixtures of hydroxylated apocarotenoid obtained by biotransformations were used directly as a substrate source for the six norisoprenoid UGTs, and the generated products were analyzed by LC-MS (Supplemental Figures S7-S12). The peak areas of the UGT products were determined in the ion traces of their pseudo molecular ions, the highest value was set to 100% and the relative quantities of the peak areas for the other UGTs calculated (Table 1). This strategy allows to determine the optimal enzyme for a substrate, but it does not allow to determine the preferred substrate for an enzyme due to the different ionizability of the substrates. MpUGT86C10, NbUGT73A25, NbUGT73A24, and NbUGT72AY1 are efficient C_13_-apocarotenoid UGTs, although NbUGT73A24 and NbUGT73A25 also transfer glucose to quercetin, kaempferol, and N-feruloyl tyramine and NbUGT72AY1 can glucosylate scopoletin (Sun et al. 2019). While MpUGT86C10 preferentially glucosylates norisoprenoids at the cyclohexene ring, as it is most active with 3-hydroxy-*α*-ionone, 4-hydroxy-*β*-ionone, 3-hydroxy-*α*-damascone, and 4-hydroxy-*β*-damascone, NbUGT73A25 promotes the transfer of glucose to the butene side chain. The latter enzyme prefers *α*-ionol, *β*-ionol, 3-hydroxy-*α*-ionol, and 4-hydroxy-*β*-ionol as substrate (Table 1). However, MpUGT86C10 is also able to glucosylate *α*- and *β*-ionol in the side chain and NbUGT73A25 can modify 4-hydroxy-*β*-ionone and 3-hydroxy-*α*-damascone at the hydroxyl group in the cyclohexene ring. NbUGT72AY1 is the most active enzyme for the glucosylation of 3-oxo-*α*-ionol (Table 1). Although kinetic data for the hydroxylated C_13_-apocarotenoids could not be determined due to a lack of purified reference material these results clearly show that MpUGT86C10 is an efficient UGT to produce a range of norisoprenoid glucosides.

### Identification of natural substrates of MpUGTs and NbUGTs

A physiologic aglycone library, which is enriched in naturally occurring aglycones can be used to reveal the natural substrates of UGTs (Bönisch et al. 2014; Sun et al. 2019). Therefore, small molecule glycosides were isolated by solid phase extraction from mint leaves and the aglycones were liberated by glycosidase treatment and analyzed by GC-MS (Figure 2). The aglycone mixture contained the C_13_-apocarotenols 3-hydroxy-*α*-damascone, 3-oxo-*α*-ionol, 3-hydroxy-7,8-dihydro-*β*-ionol, and 3-oxo-7,8-dihydro-*α*-ionol (Blumenol C), which have not been isolated from mint plant before, while in the mint glycoside extract the glucosides of these norisoprenoids could be tentatively identified by LC-MS (Figure 2). The same glucosides were produced when the aglycone library was used as substrate source for MpUGT86C10, while no glucoside was generated by the empty vector control (Figure 2 and Supplemental Figure S13). Therefore, the natural substrates of MpUGT86C10 in *M. × piperita* appear to be 3-hydroxy-*α*-damascone, 3-oxo-*α*-ionol, 3-hydroxy-7,8-dihydro-*β*-ionol and 3-oxo-7,8-dihydro-*α*-ionol. Similarly, glycosides were isolated from *N. tabacum* leaves and fractionated into 30 fractions by preparative LC. *N. tabacum* was used for the isolation of glycosides because tobacco is readily available in large quantities. The aglycones were liberated in the fractions using a glycosidase (Sun et al. 2019). 3-Oxo-*α*-ionol and 3-oxo-7,8-dihydro-*α*-ionol (Blumenol C) could be detected by GC-MS in fractions 9-11 (Supplemental Figure S14), and the corresponding glucosides provisionally detected in the tobacco fractions by LC-MS. The 3-oxo-*α*-ionyl glucoside was successfully reconstituted from the aglycone library by NbUGT72AY1 and confirmed the preference of this enzyme for 3-oxo-α-ionol already seen in Table 1 (Supplemental Figure S14). Further UGTs were not tested with the aglycone library.

**Fig. 2.**
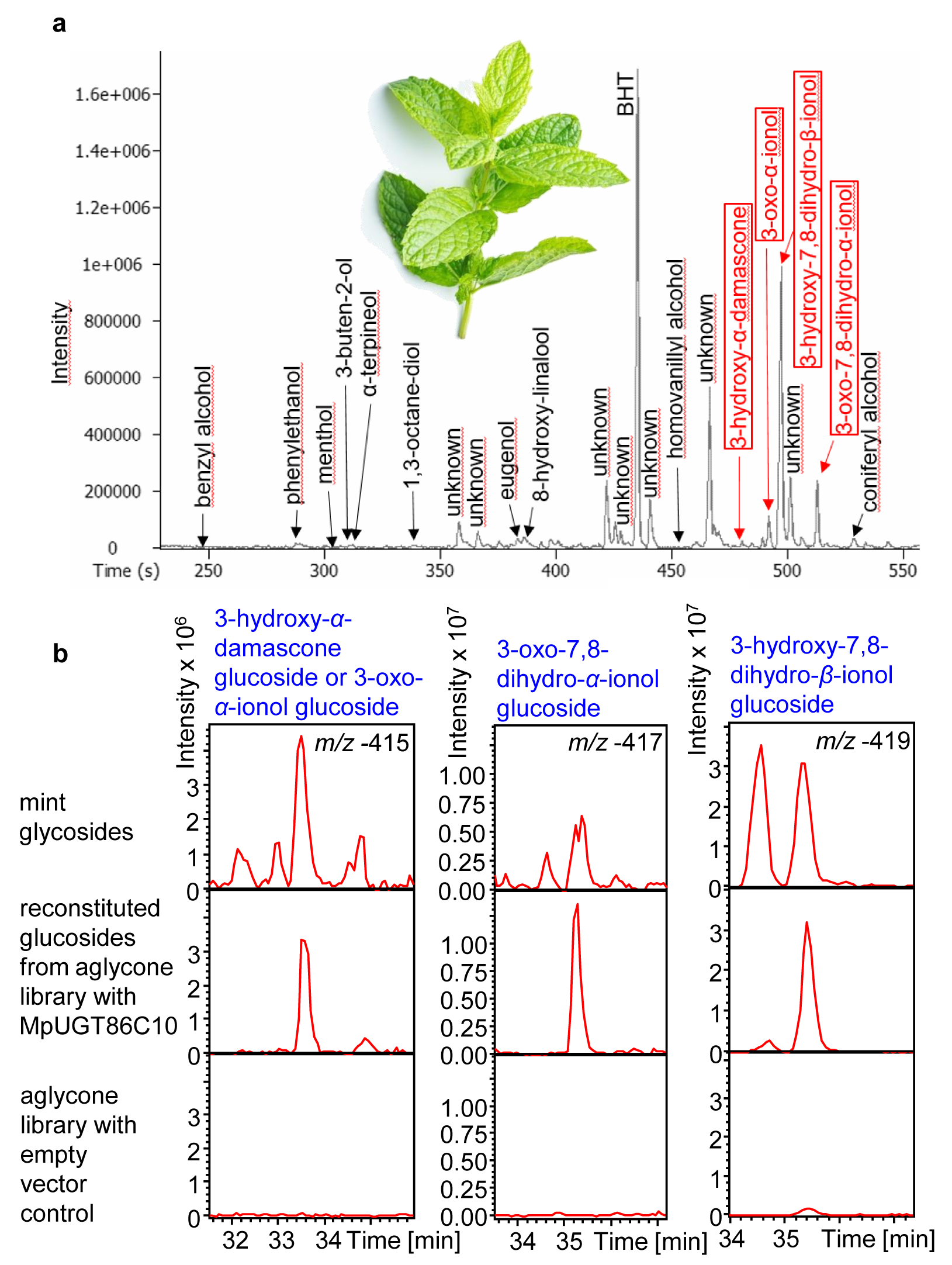
A mint aglycone library was used as substrate source for recombinant MpUGT86C10 from *M. × piperita*. (**a**) Volatile metabolites released by glucosidase (Rapidase) from a mint glycoside extract (aglycone library) were analyzed by GC-MS. C_13_-apocarotenoids are shown in red boxes. BHT butylated hydroxytoluene stabilizer. (**b**) The aglycone library was subsequently employed as substrates for MpUGT86C10. The mint glycoside extract and the aglycone library incubated with empty vector control served as positive and negative control, respectively. Diagnostic ions are explained in Supplemental Table S1.

### Agroinfiltration of *N. benthamiana* for transient expression of C_13_-apocarotenol *UGT* genes and demonstration of UGT *in planta* activity

To investigate the *in planta* function of the norisoprenoid UGTs all UGTs except for NbUGT85A73, which showed lower activity for the C_13_-apocarotenols, were transiently overexpressed in *N. benthamiana* leaves using an established viral vector system (Sun et al. 2019). As controls, an empty vector (CO) was infiltrated and untreated wild type leaves (WT) were used. *N. benthamina* leaves were treated with *Agrobacterium* from abaxial and were harvested 7 and 10 days after agroinfiltration. During this period, the leaf color changed from green to light yellow (Figure 3). To test whether the infiltrated genes were successfully expressed in the leaves, RNA and proteins were extracted to perform qRT-PCR and enzyme activity assays, respectively. Transcript levels of the infiltrated genes were significantly increased in the agroinfiltrated leaves in comparison with CO and WT samples (Figure 3). Protein extracts obtained from *UGT72AY1-, UGT73A25-, UGT73A24-, UGT709C6-, and UGT86C10-*infiltrated leaves 7 days post-infiltration were incubated with *α*-ionol and *β*-ionol as acceptor substrates and UDP-glucose as donor. All samples produced glucosides except for *UGT709C6*, WT and CO protein extracts (Figure 3). The highest level of *α*- and *β*-ionyl glucoside was produced by UGT73A24 and UGT86C10. Therefore, the infiltrated genes were successfully overexpressed in *N. benthamiana* leaves and contribute to the formation of ionyl glucosides *in vitro*. To detect changed metabolite levels in agroinfiltrated leaves, we performed targeted LC-MS analysis with extracts obtained from *UGT72AY1-, UGT73A25-, UGT73A24-, UGT709C6-, and UGT86C10-*leaves and with samples from CO and WT leaves. The levels of the major *Nicotiana* C_13_-apocarotenyl glucosides, 3-oxo-*α*-ionyl glucoside and 3-oxo-7,8-dihydro-*α*-ionyl glucosides (Cai et al. 2014) were not significantly increased in the agroinfiltrated leaves in comparison with CO and WT leaves.

**Fig. 3.**
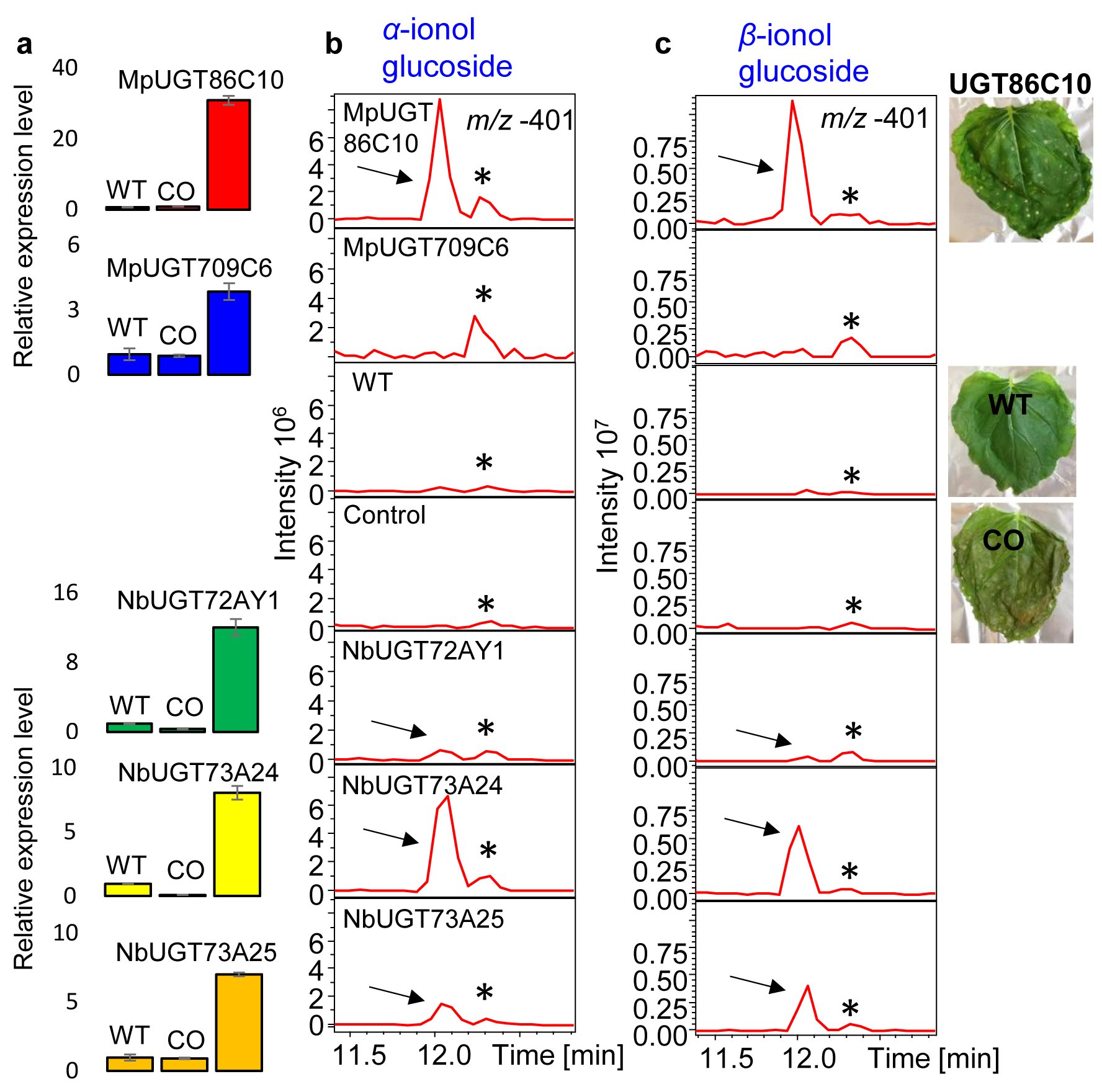
Agroinfiltration of *NbUGT72AY1, NbUGT73A24, NbUGT73A25, MpUGT86C10* and *MpUGT709C6* in *N. benthamiana*. (**a**) QPCR analysis was performed with WT, CO and agroinfiltrated leaves (*NbUGT72AY1, NbUGT73A24, NbUGT73A25, MpUGT86C10* and *MpUGT709C6*). Tobacco UGTs were analyzed at 7 d and mint UGTs at 10 d post-infiltration. Triplicates were analyzed. (**b**) Products obtained by enzyme assays with protein extracts from the WT, empty vector control, *NbUGT72AY1-, NbUGT73A24-, NbUGT73A25-, MpUGT86C10-* and Mp*UGT709C6*-infiltrated leaves and the substrates *α*-ionol were analyzed by LC-MS. *MpUGT709C6* and *MpUGT86C10* extracts were incubated at 30 °C for 2 h, all other extracts were incubated at 30 °C for 1 h. Ionyl glucoside (indicated by a black arrow) was detected at ion trace *m/z* 401 [M+HCOO]^-^ and verified by MS2 analysis. A co-eluting plant metabolite is indicated by a black asterisk. (**c**) Same as in (**b**) but *β*-ionol was used as substrate. Phenotypes of WT, CO and *UGT86C10*-infiltrated leaves are shown on the right side.

Due to these ambiguous results, we assumed that limited substrate availability prevents glucoside formation. Therefore, we decided to demonstrate the C_13_-apocarotenoid UGT activity of MpUGT86C10 by agroinfiltration of the corresponding gene in *N. benthamiana* and concurrent infiltration of *α*- and *β*-ionol to counteract limited substrate availability. As controls, an empty vector was agroinfiltrated (CO) and untreated wild type leaves (WT) were used. *N. benthamina* leaves were infiltrated with *MpUGT86C10* from abaxial and after 3 days, leaves were dripped with *α*- and *β*-ionol from adaxial using a pipette. All samples were harvested 5 and 7 days after co-infiltration of the C_13_-apocarotenol substrates. To test whether MpUGT86C10 was successfully expressed in the leaves, proteins were extracted and enzyme activity assays were performed (Supplemental Figures S15-S17). *MpUGT86C10*-infiltrated samples (Supplemental Figure S17) produced high amounts of ionyl glucosides in particular *α*-ionyl glucoside *in vitro*, while protein extracts from WT and CO plants yielded only minor quantities. The results indicated that *MpUGT86C10* was successfully expressed in *N. benthamiana* leaves and catalytically active protein was produced.

To quantify ionyl glucoside concentrations in the co-infiltrated leaves, we also performed targeted LC-MS analysis of methanolic extracts obtained from *MpUGT86C10-*infiltrated, CO, and WT leaves and corresponding leaves co-infiltrated with *α*- and *β*-ionol (Figure 4). Low quantities of *α*- and *β*-ionyl glucoside were found when WT and CO leaves were infiltrated with the ionols, confirming that *N. benthamiana* plants express endogenous C_13_-apocarotenol UGTs in their leaves. The level of *α*-ionyl glucoside was significantly and that of *β*-ionyl glucoside slightly increased in the *MpUGT86C10-*infiltrated leaves, which were co-infiltrated with the ionols in comparison to WT and CO leaves. In addition, we putatively identified 7,8-dihydro-*α*- and 7,8-dihydro-*β*-ionyl glucoside (Figure 4) after co-infiltration of *MpUGT86C10*. The detection of 7,8-dihydro-ionols confirmed the already described 7,8-dehydrogenase activity of C_13_-apocarotenoids in *Nicotiana* (Tang and Suga 1994). Thus, the result clearly shows that MpUGT86C10 acts as C_13_-apocarotenol UGT *in planta*.

**Fig. 4.**
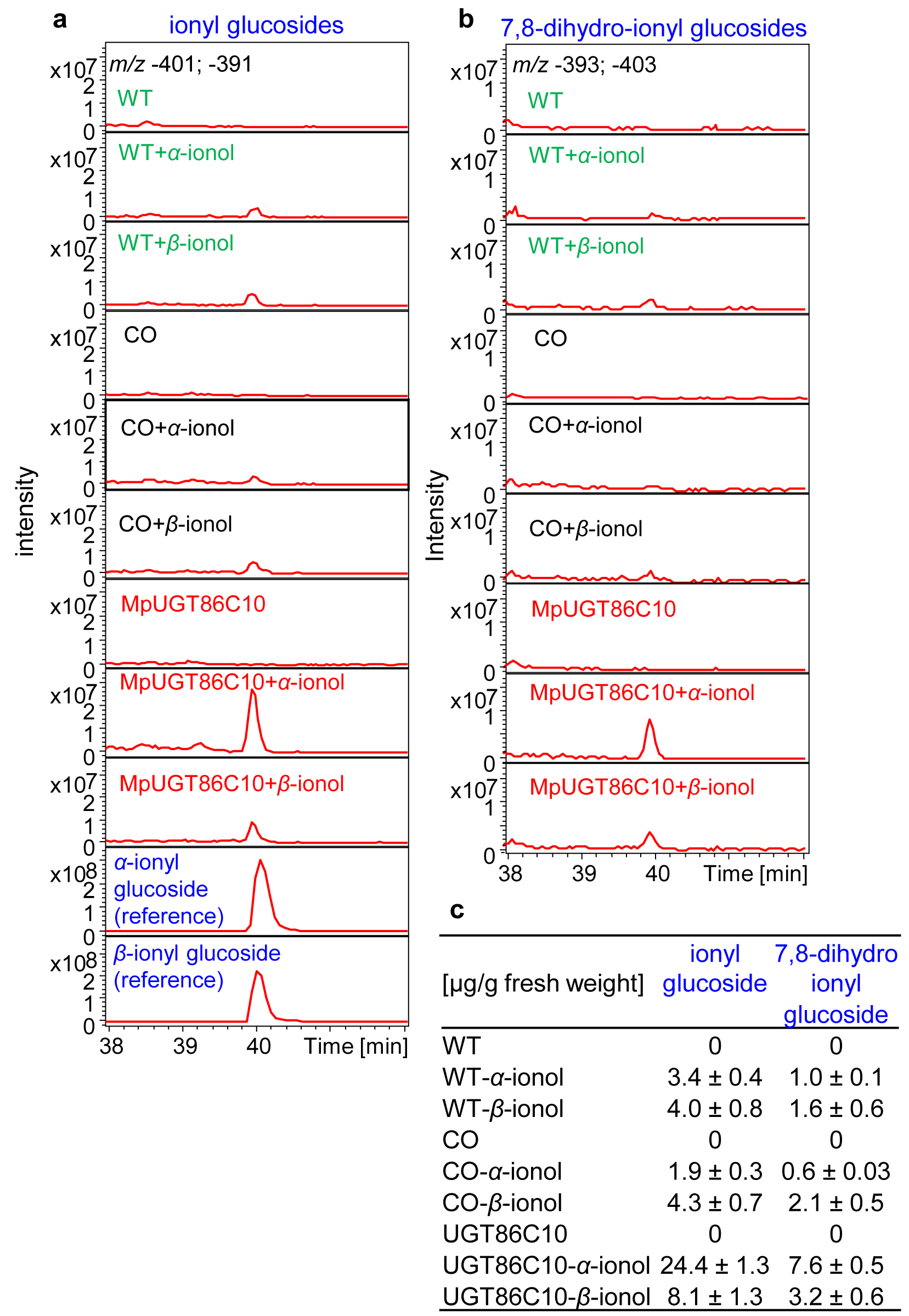
Co-infiltration of *MpUGT86C10* and potential substrates (*α*- and *β*-ionol) into *N. benthamiana* leaves. (**a**) LC-MS analysis of extracts obtained from infiltrated leaves to detect ionyl glucosides at *m/z* -401 and -391. Untreated wild type plants WT, plants infiltrated with an empty vector CO, and infiltrated with *UGT86C10*. (**b**) Detection of 7,8-dihydro-ionyl glucoside at *m/z* -393 and -403. (**c**) Mean concentration (µg/g fresh weight) of triplicate measurements.

### C_13_-apocarotenols reduce germination rates and embryo size of *N. benthamiana* seeds

To further clarify the *in planta* role of C_13_-apocarotenyl glucosides, we carried out a germination test since norisoprenoid glucosides may have allelopathic activity. *N. benthamiana* seeds were subjected to different concentrations (0.1; 1; 10 and 100 mM) of *α*-ionol and *α*-ionyl glucoside, respectively (Figure 5). While the seeds germinated readily at 0.1 and 1 mM *α*-ionol, 10 mM and 100 mM of the isoprenoid reduced the germination rate (68% and 65%, respectively) and the overall embryo size significantly, and resulted in a lighter leaf color. The effect of *α-*ionyl glucoside was even more pronounced. A 10 mM solution already inhibited the germination, completely. Consequently, we conclude that α-ionyl glucoside has a stronger effect on seed germination and is potentially more phytotoxic than its aglycone *α*-ionol. Although the *Nicotiana*/ionol combination is a model system, the results suggest that possible biological functions of the glucosides should not be neglected and the thesis that glycosylation leads to detoxification has no general validity.

**Fig. 5.**
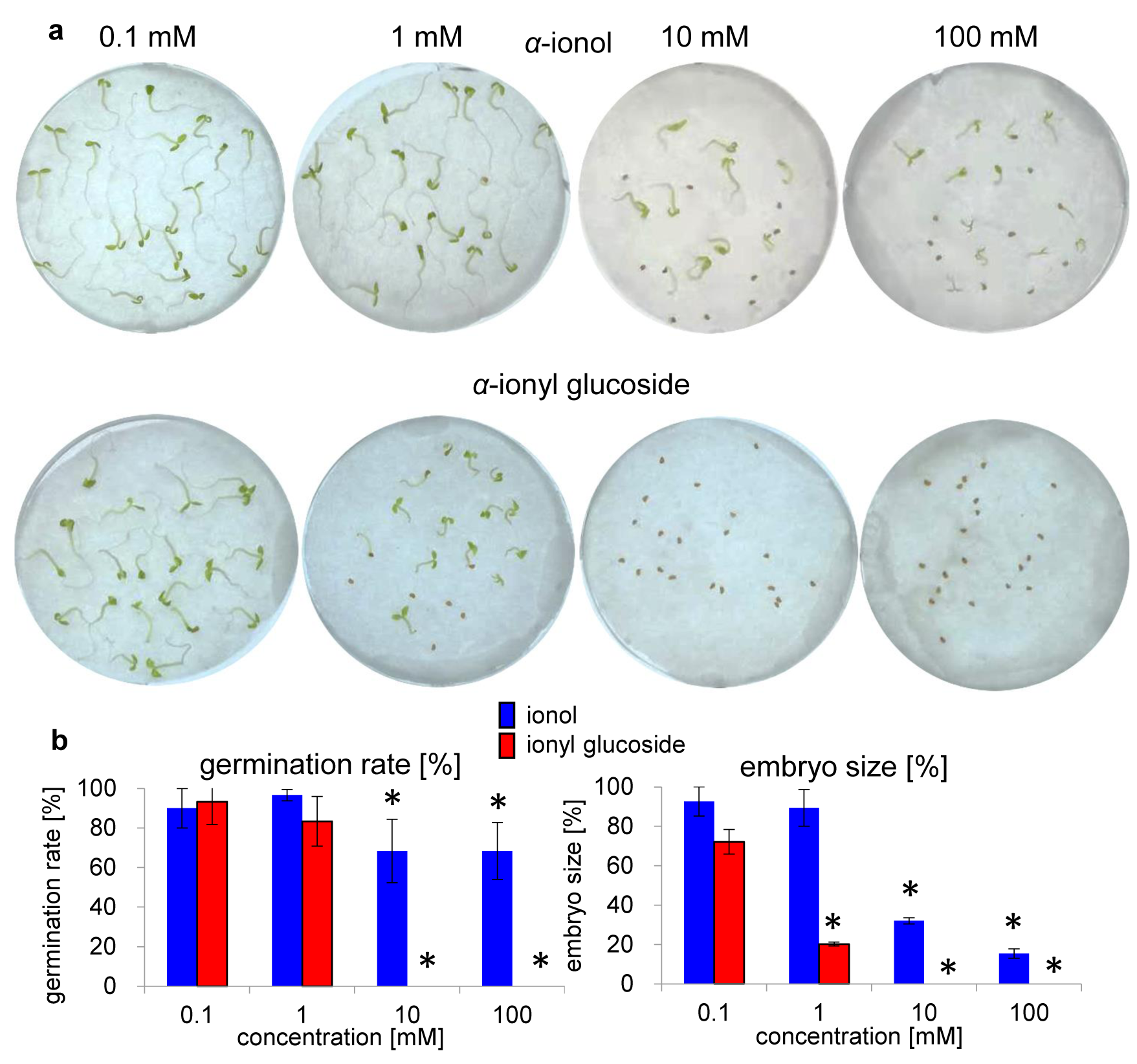
Germination of *N. benthamiana* seeds in the presence of C_13_-apocarotenols. (**a**) Effect of 0.1-100 mM *α*-ionol and *α*-ionyl glucoside on *N. benthamiana* seedlings. Triplicates of 20 seeds per plate were analyzed. (**b**) Germination rate and embryo size in % after application of 0.1-100 mM of *α*-ionol and *α*-ionyl glucoside. Data are presented as mean ± SE of three repetitions (twenty seeds per treatment). Asterisks indicate significant differences in comparison with the lowest concentration (Student’s t-test: *P < 0.05).

## Discussion

Although the production and function of apocarotenoids has attracted much attention in recent years, their metabolism, including hydroxylation and glycosylation, has rarely been studied, with the exception of the plant hormone ABA (Finkelstein 2013).

### Identification of unprecendented C_13_-apocarotenol UGTs

Since no C_13_-apocartenol UGT has been biochemically characterized to date, we used commercially available *α*- and *β*-ionol as model norisoprenoids to screen tobacco UGTs that were recently characterized (Sun et al. 2019). In addition, recombinant *M. × piperita* UGTs whose genes were derived from a mint leaf transcriptome database (http://langelabtools.wsu.edu/mgr/home), were used as candidates (Vining et al. 2017). UGTs from mint have not yet been analyzed. We focused on UGTs strongly expressed in leaves, as vegetative tissue is a rich source of norisoprenoid glycosides (He et al. 2015; Winterhalter and Rouseff 2002; Wirth et al. 2001). Out of 16 plant UGTs analyzed in this study (ten from *N. benthamiana* and 6 from *M. x piperita*), six proteins from distinct UGT families (Supplemental Figure S5) accepted *α*- and *β*-ionol as acceptor substrate, exhibiting apparent *K*_M_ values of 6 to 131 µM (Table 2). In comparison, UGT75L6 from *G. jasminoides*, which forms a glucose ester showed an apparent *K*_M_ value for the C_20_-apocarotenoid crocetin of 0.46 mM (Nagatoshi et al. 2012), while the apparent *K*_M_ value for the C_10_-apocarotenoid 3-hydroxy-*β*-cyclocitral of UGT709G1 from *C. sativus* was 64.0 µM (Diretto et al. 2019). UGTs acting on terpenes show *K*_M_ values from 9-463 µM for monoterpenols such as citronellol, geraniol, nerol and linalool and kcat/*K*_M_ values from 40-2600 M^-1^ sec^-1^ for the same substrates (Bönisch et al. 2014; Wu et al. 2019). Since *α*- and *β*-ionol have not yet been found as aglycones in plants, further hydroxylated C_13_-apocarotenols were synthesized. Therefore, 3-hydroxy-*α*- and 4-hydroxy-*β*-ionol, 3-hydroxy-*α*- and 4-hydroxy-*β*-ionone, as well as 3-hydroxy-*α*- and 4-hydroxy-*β*-damascone were produced by P450-catalyzed whole cell biotransformation of *α*/*β*-ionols, *α*/*β*-ionones, and *α*/*β*-damascones. The glycosides of 3-hydroxy-*α*-ionol and 4-hydroxy-*β*-ionone have been isolated from stinging nettle (*Urtica dioica* L.) leaves (Neugebauer et al. 1995) and raspberry fruits (*Rubus idaeus*), (Pabst et al. 1992b) respectively. The reaction products were used as acceptor substrates for the UGTs and the relative activities calculated (Table 1). UGTs could be distinguished according to their selectivities towards the glucosylation of the hydroxyl group attached to the cyclohexene ring (MpUGT86C10) or the butene side chain (NbUGT73A25). Both types of norisoprenoid glycosides have been isolated from plant sources, 3-hydroxy-*α*-ionol 9-*O*-*β*-D-apiofuranosyl-(1-6)-*β*-D-glucopyranoside (side chain attachment) and 4-hydroxy-*β*-ionone 4-*O*-*α*-L-arabinofuranosyl-(1-6)-*β*-D-glucopyranoside (ring attachment) from *Cydonia vulgaris* (Tommasi et al. 1996) and raspberry (Pabst et al. 1992b), respectively. The result confirmed the validity of our strategy. The peculiarity of MpUGT86C10 is illustrated by the protein sequence analysis and catalytic specificity. This biocatalyst is separated from the other sequences in the phylogenetic tree (Supplemental Figure S5) and shows a preference for ring glucosylation (Table 1). The amino acid sequence analysis does not yet allow identifying the amino acids responsible for this particular characteristic. Although successful overexpression of the agroinfiltrated UGT genes in *N. benthamiana* was demonstrated by qPCR and enzyme activity assays (Figure 3), except for MpUGT709C6, targeted LC-MS analysis of norisoprenoid glycosides was inconclusive probably due to limited substrate availability. In other words, there is probably not enough free apocarotenoid available in the leaves to detect a significant increase in glycoside formation after agroinfiltration. Therefore, *α*-ionol and *β*-ionol were co-infiltrated after agroinfiltration (Figure 4). After addition of the ionols, the concentration of their corresponding glucosides increased significantly in *MpUGT86C10-*infiltrated leaves proving the C_13_-apocarotenol UGT activity of MpUGT86C10 but also revealing endogenous *α*-ionol and *β*-ionol UGT activity in *N. benthamiana* leaves. The putative identification of the hexosides of 7,8-dihydro-*α*-ionol and 7,8-dihydro-*β*-ionol is supported by the detection of a 7,8-dehydrogenase in *N. tabacum* (Tang and Suga 1994). Studies on gene function analysis using agroinfiltration of candidate genes in combination with metabolite analysis always have to reckon with limited substrate availability, which is a major disadvantage of this strategy.

### C_13_-apocarotenoid UGTs are promiscuous biocatalysts with different functions

UGTs of five different classes (72, 73, 85, 86, and 709) were able to transfer a sugar molecule onto C_13_-apocarotenoids. The enzymes NbUGT72AY1, NbUGT73A24, NbUGT73A25, and NbUGT85A73 have recently been shown to be promiscuous enzymes. They glucosylated 18, 17, 17, and 19 small molecules out of a selection of 27 different metabolites, which were known to occur naturally in *Nicotiana* species including scopoletin, benzyl alcohol, 2-phenylethanol, kaempferol, and 3-cis-hexenol, and structurally related aliphatic, branched chain, and phenolic metabolites (Sun et al. 2019).

Further studies demonstrated that members of the UGT72 family glucosylate flavonols (Yin et al. 2017), flavanones, anthocyanins (Zhao et al. 2017), and monolignols (Lanot et al. 2008) and are most probably involved in lignin biosynthesis (Wang et al. 2012). UGT73 family members catalyze the 3-*O*-glucosylation of the sapogenins oleanolic acid and hederagenin (Augustin et al. 2012; Erthmann et al. 2018) as well as the conversion of brassinosteroid phytohormones (Poppenberger et al. 2005). UGT73 enzymes are also responsible for the 7-*O*-glucosylation of kaempferol and quercetin 3-*O*-rhamnoside (Jones et al. 2003), the transformation of salicylic acid and scopoletin (Simon et al. 2014) as well as feruloyl tyramine (Sun et al. 2019). Therefore, UGT72 and 73 members from *N. benthamiana* show broad substrate tolerance and use phenolic compounds, including flavonols, but also aliphatic alcohols such as terpenoids as acceptor molecules.

UGT85A73 is probably implicated in the glucosylation of tobacco flower volatiles as it is strongly expressed in tobacco blossom and glucosylates nonpolar, low-molecular-weight compounds (Caputi et al. 2012; Sun et al. 2019). UGT85s from peach, grape, tea plant, and kiwifruit catalyze the glucosylation of aliphatic alcohols including linalool, geraniol, citronellol, hexanol, (Z)-3-hexenol, octanol and also volatiles such as 2-phenylethanol, benzyl alcohol, and furaneol (Bönisch et al. 2014; Jing et al. 2019; Song et al. 2018; Wu et al. 2019).

The related UGT709C2 protein shows catalytic activity towards 7-deoxyloganetate, a precursor of loganin and secologanin (Asada et al. 2013). Interestingly, UGT709G1 has only recently been identified as a novel C_10_-apocarotenoid UGT from saffron (*C. sativus*) that produces picrocrocin, the precursor of safranal from 3-hydroxy-*β*-cyclocitral (Diretto et al. 2019). Strikingly, picrocrocin was found in *N. benthamiana* when 3-hydroxy-*β*-cyclocitral was provided by CCD2, indicating apocarotenoid UGT activity in tobacco (Diretto et al. 2019). Although UGT86C4 (Bhat et al. 2013), UGT86C1 (Ono et al. 2006), and UGT86C3 (Noguchi et al. 2008) have been isolated from different plants, no biochemical analyses have been performed to date.

### Aglycone libraries are versatile tools for enzyme function analysis

Physiologic libraries of aglycones have proven to be versatile tools for identifying natural acceptor substrates of UGTs especially if potential metabolites are not commercially available (Bönisch et al. 2014). The broad substrate compatibility of UGTs requires a broad spectrum of potential aglycones to be tested, which is often hindered by a biased collection of acceptor molecules. This difficulty has recently been alleviated by the use of aglycone libraries to identify *in planta* substrates (Bönisch et al. 2014). The powerful tool was used to identify unknown substrates of recombinant plant UGTs. Aglycone libraries were easily prepared by isolation and subsequent glycosidase treatment of glycoconjugates. The natural glycosides were then reconstituted by enzyme activity assays. This approach allowed the identification of UDP-glucose:monoterpenol UGTs from grapevine (Bönisch et al. 2014) and two feruloyl tyramine UGTs from *N. benthamina* (Sun et al. 2019) and identified kaempferol, quercetin, abscisic acid and three unknown natural metabolites as putative *in planta* substrates of UGT71 family members (Song et al. 2015). In this study, a collection of aglycones liberated from glycosides isolated from *M. × piperita* and *N. benthamiana* enabled the characterization of the first C_13_-apocarotenoid UGTs (Figure 2 and Supplemental Figures S13 and S14). MpUGT86C10 readily glucosylated 3-hydroxy-*α*-damascone, 3-oxo-*α*-ionol, 3-oxo-7,8-dihydro-*α*-ionol, and 3-hydroxy-7,8-dihydro-*β*-ionol (Blumenol C), metabolites previously unknown in mint. We assume that other members of the UGT86 family, of which no enzyme has yet been biochemically characterized, are also able to glucosylate apocarotenoids. The norisoprenoids found in mint have frequently been detected in different plant tissues (Winterhalter and Rouseff 2002) and are produced by cell suspension cultures of *V. vinifera* from *β*-ionone and dehydrovomifoliol (Mathieu et al. 2009). Studies with carotenoid hydroxylase mutants suggested that C_13_-apocarotenoids derive from xanthophylls rather than through secondary oxygenation of *α*- and *β*-ionone (Lätari et al. 2015; Mathieu et al. 2009). However, reduction of the 7,8 double bond appears to take place after the formation of the C_13_-structure (Tang and Suga 1994), which was also confirmed by our results. Some fruits and flowers contain high amounts and a hugh diversity of apocarotenoid glycosides (Winterhalter and Rouseff 2002), which indicates increased xanthophyll degradation during ripening or senescence of the fruits and changes in the use of xanthophyll precursors and secondary modifications (Lätari et al. 2015).

### Possible biological functions of C_13_-apocarotenoid UGTs

Lately, carotenoid and apocarotenoid metabolism in *Arabidopsis thaliana* was investigated in response to enhanced carotenoid production upon phytoene synthase overexpression (Lätari et al. 2015; Zhou et al. 2015). Although overexpression of the carotenoid biosynthesis gene led to a dramatic accumulation of mainly *β*-carotene in non-green tissues, carotenoid levels remained unchanged in leaves. In green tissues, the increased pathway flux was compensated by generation of a high level of C_13_-apocarotenoid glycosides, including 3-oxo-*α*-ionyl, 3-hydroxy-5,6,-epoxy-*β*-ionone, and 6-hydroxy-3-oxo-*α*-ionone glycoside (Lätari et al. 2015). In contrast to leaves, apocarotenoid glycosides were absent in the roots.

Similarly, green tissues of phytoene synthase-overexpressing tomato and *N. tabacum* plants also showed only slightly increased carotenoid concentrations compared with wild-type plants (Busch et al. 2002; Fray et al. 1995). The authors proposed that tissue-specific capacities to synthesize xanthophylls determine the modes of carotenoid accumulation and apocarotenoid generation. The multiple functions of carotenoids and apocarotenoids such as ABA require regulation of their synthesis and the production, release, transformation and disposal of their breakdown products. Similarly, glycosylation of apocarotenoids catalyzed by NbUGT72AY1, NbUGT73A24, NbUGT73A25, and MpUGT86C10 may function as a valve adjusting carotenoid and apocarotenoid steady-state levels in leaves.

One of the functions of glycosylation is the detoxification of metabolites through increased solubility to enable cell transport and vacuolar sequestration (Song et al. 2018). Electrophilic *α,β*-unsaturated carbonyls can cause various detrimental effects due to interactions with proteins, and these functional chemical groups are frequently found in carotenoid degradation products. In order to be prepared for the detoxification of different metabolites, it is advisable to be able to glucosylate a wide range of alcohols. Therefore, apocarotenol UGTs show substrate promiscuity similar to other UGTs of the secondary metabolism (Song et al. 2018). The C_13_-apocarotenoid glycosides also perform vital functions in plants since crosses of *phytoene synthase* overexpessing lines with *carotenoid cleavage dioxygenase* deficient mutants produced lethal seedlings suggesting deleterious effects at high fluxes into the carotenoid pathway when the detoxification mechanism into apocarotenoid glycosides is dysfunctional (Lätari et al. 2015). Therefore, we propose a biosynthetic pathway for the production of 3-hydroxy-7,8-dihydro-β-ionyl-, 3-oxo-α-ionyl-, and 3-oxo-7,8-dihydro-α-ionyl glucoside in *M. × piperita* and *N. benthamiana* (Figure 6) in accordance to a published scheme for *A. thaliana* (Lätari et al. 2015).

**Fig. 6.**
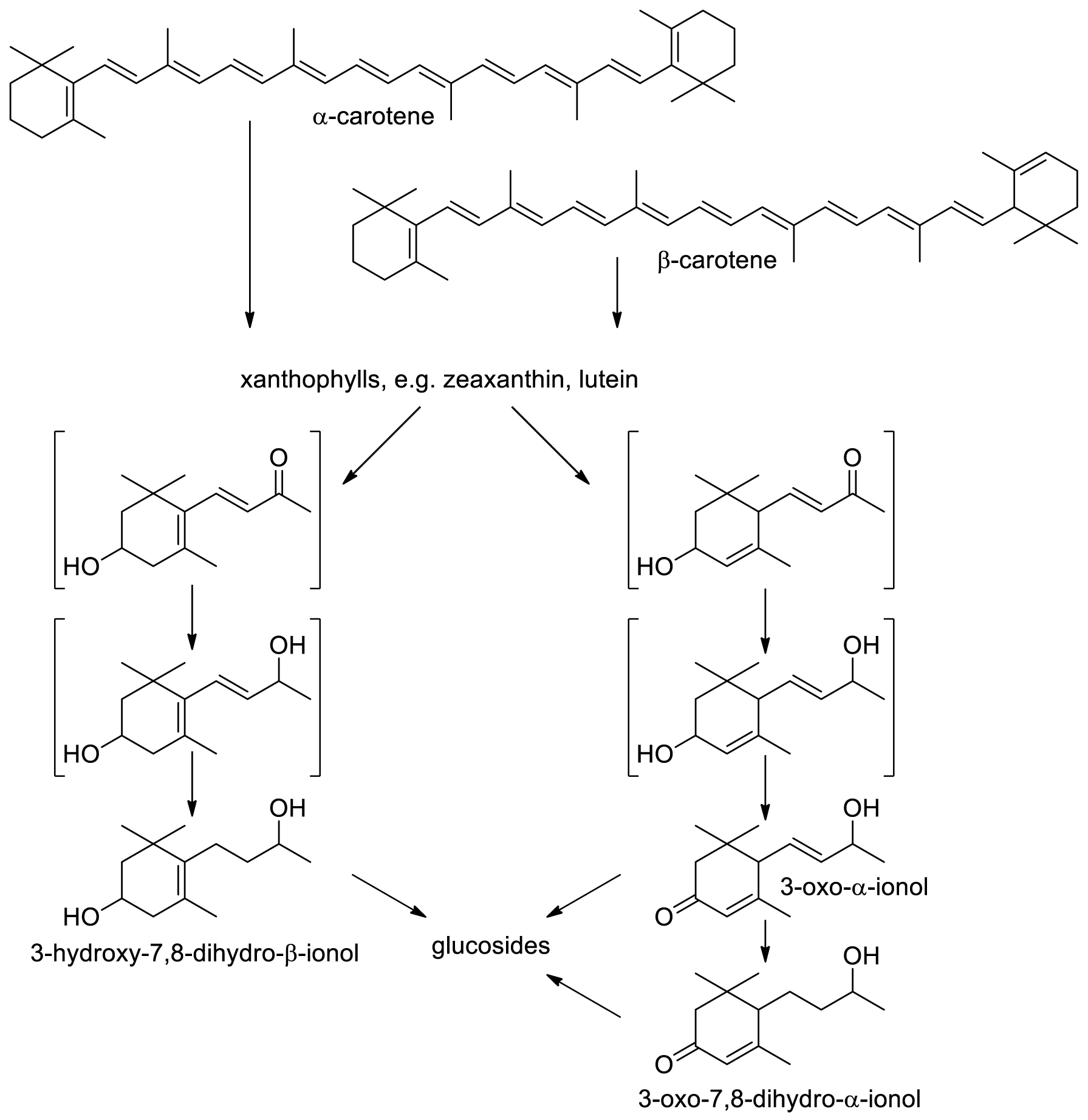
Formation of apocarotenyl glucosides in *M. × piperita* and *N. benthamiana.*

### C_13_-apocarotenol glucosides could act as allelochemicals and are biomarkers for colonization with arbuscular mycorrhizal fungi

Numerous allelochemicals have been isolated from a variety of plant species. They comprise different chemical families including phenolics, terpenoids, alkaloids, and other nitrogen-containing chemicals (Kong et al. 2019). Allelochemicals are released into the environment and are important in regulating the interaction between plants and other organisms. Since C_13_-apocarotenoids inhibit seed germination and impair root and shoot growth at concentrations as low as 1 mM (D’Abrosca et al. 2004; Kato-Noguchi et al. 2010; Kobayashi and Kato-Noguchi 2015) we studied the effect of glucosylation on the allelopathic activity of the model norisoprenoid α-ionol. The concentrations required for 50% growth inhibition on root and shoot growth of different plant species were 2.7–19.7 μM for 3-hydroxy-β-ionone, and 2.1–34.5 μM for 3-oxo-α-ionol (Kato-Noguchi et al. 2010). In a germination test, we could show that glucosylated *α*-ionol had an even more drastic negative effect on germination rate and embryo size compared to unbound *α*-ionol, which showed much weaker allelopathic activity than 3-hydroxy-β-ionone and 3-oxo-α-ionol (Figure 5). Although C_13_-apocarotenyl glucosides have often been identified together with their allelopathic aglycones in plants, nothing is known about the phytotoxic potential of the glucosides. (Dietz and Winterhalter 1996; Llanos et al. 2010). Our results show that glucosylation does not always imply detoxification and consequently a reduction of the phytotoxic effect of the aglycones, but can also enhance it. Similarly, saponins (steroid glucosides) are more phytotoxic than their aglycones where the allelopathic activity is attributed to their surfactant character (Ghimire et al. 2019; Mushtaq and Siddiqui 2018). In addition, phytotoxic iridoid glucosides and flavonoid glycosides were found in roots of *Vesbascum thapsus* (Pardo et al. 1998) and leaves of *Myrcia tomentosa* (Imatomi et al. 2013), respectively.

Glycosylation is thought to be a prerequisite for the transport of secondary metabolites in plants (Yazaki et al. 2008) and therefore might also facilitate exudation of the allelochemicals by roots (Weston et al. 2012). Glycosylation increases the mobility of plant metabolites in soil as was shown for flavone *O*-glycosides isolated from allelopathic rice seedlings (Kong et al. 2007).

Only recently, C_13_-apocarotenyl glucosides (11-hydroxyblumenol C-9-*O*-, and 11-carboxyblumenol C-9-*O*-glucoside) have been proposed as shoot markers of root symbiosis with arbuscular mycorrhizal fungi (AMF) (Wang et al. 2018). Blumenol-(3-oxo-α-ionol derivatives)-type metabolites accumulate in roots of different plants after AMF inoculation and their concentration is highly correlated with the fungal colonization rate (Strack and Fester 2006). Glycosylation of the blumenols usually occurs at the hydroxyl group at the C9 position (side chain). The glycosyl moiety can be a monosaccharide or combinations of different sugars (Wang et al. 2018). The type of decorations is highly species-specific. Some of these glucosides are formed in the roots and transported to the shoots (Wang et al. 2018). Future studies should analyse whether MpUGT86A10 related UGTs are involved in the glucosylation of blumenols, which are formed when plant roots are colonized by AMF. *N. benthamiana* and *M. x piperita* root tissue show high expression levels of the C_13_-apocartenol *UGT* genes in comparison to other *UGTs* (Supplemental Figure S2).

In this study, unprecedented C_13_-apocarotenol UGTs were characterized, natural substrates of UGT86 class members were identified, and the versatility of aglycone libraries to reveal *in planta* substrates was demonstrated. In the future, the results will shed more light on the importance of apocarotenoid glycosides in plants, animals and human, and will enable further studies to investigate their role as signalling substances.

## Materials and Methods

### Chemicals and plant materials

Chemicals and solvents were purchased from Sigma (Steinheim, Germany), Aldrich (Steinheim, Germany), Roth (Karlsruhe, Germany), and Fluka (München, Germany). Uridine 5-diphosphate (UDP), UDP-glucose, *α*-ionol and *β*-ionol (≥ 90%) were purchased in analytical grade from Sigma-Aldrich. Peppermint (*M. × piperita*) and tobacco plants (*N. benthamiana*) used for the isolation of the UGT genes were cultured at room temperature in a growth chamber maintained at 22 ± 2 °C with a 16 h light, 8 h dark photoperiod and a light intensity of 70 ± 10 μmol m^-2^ sec^-1^, respectively. For over-expression and molecular analyses, tobacco leaves were injected with viral vectors (Sun et al. 2019) and harvested 7 and 10 days after agroinfiltration.

### Selection of UGTs – UGTs from *Mentha × piperita*

To find putative UGTs in *M. × piperita* (*MpUGTs*), a database search was performed using the transcriptome data from the Mint Genomics Resource (MGR) at Washington State University (http://langelabtools.wsu.edu/mgr/home) since the draft genome sequence was not yet available. Three full-length putative *MpUGT* nucleotide sequences were assembled based on overlapping expressed sequence tags. The genes were designated *MpUGT86C10, MpUGT708M1*, and *MpUGT709C6* (https://prime.vetmed.wsu.edu/resources/udp-glucuronsyltransferase-homepage). The allelic forms *MpUGT708M2* and MpUGT709C7/8 were obtained during the isolation of *MpUGT708M1* and *MpUGT709C6*, respectively.

### Selection of UGTs – UGTs from *Nicotiana benthamina*

Ten UGTs were recently isolated from *N. benthamina* (NbUGTs) and designated *UGT71AJ1, UGT72AX1, UGT72AY1, UGT72B34, UGT72B35, UGT73A24, UGT73A25, UGT85A73, UGT85A74*, and *UGT709Q1* (Sun et al. 2019). Nucleotide and amino acid sequence analyses were performed with the Geneious Pro 5.5.6 software (Biomatters, http://www.geneious.com/).

### Determination of gene expression levels and cloning of plasmid constructs

Total RNAs were isolated from *M. × piperita* and *N. benthamiana* leaves using the Rneasy plant mini kit (Qiagen, Hilden, Germany) and CTAB extraction, respectively (Sun et al. 2019), followed by DNase I (Fermentas, St. Leon-Rot, Germany) treatment and reverse transcription to prepare cDNA. The transcribed cDNA was used as template for the PCR reactions, which were carried out in a 30 μl total reaction volume. The program was 2 min at 98 °C, one cycle; denaturation 30 sec at 98 °C, annealing 30 sec at 55 °C, elongation 1 min at 72 °C, 35 cycles; extension 10 min at 72 °C, one cycle, final temperature 8 °C, using appropriate primers (Supplemental Table S4). After gel extraction of the DNA fragments with the PCR Clean-up Gel Extraction Kit (Macherey-Nagel, Düren, Germany), the DNA fragments and the vector DNA were digested by the same restriction enzymes (Supplemental Table S4) and were ligated into pGEX-4T-1 vector. The recombinant plasmids (pGEX-4T1-UGTs) were transformed into *E. coli* NEB 10 beta. After colony PCR and restriction enzyme digestion analysis, the positive plasmids were sequenced and stored as cryostock cultures at −80 °C.

### Heterologous protein expression

Protein expression was performed with *E. coli* BL21 (DE3) pLysS containing the pGEX-4T-1 vector and the UGT sequences according to (Sun et al. 2019). Recombinant proteins were analyzed by SDS-PAGE stained with Coomassie brilliant blue R-250 (Supporting figures: Figure S3) and Western blot using anti-GST antibody and goat anti-mouse IgG fused to alkaline phosphatase. Proteins were quantified by Bradford assay.

### Substrate screening by LC-MS

To identify functional C_13_-apocarotenol UGTs, the recombinant proteins were assayed with the model norisoprenoids *α*- and *β*-ionol, and the reference material 3-oxo-*α*-ionol, which was kindly provided by Ziya Y. Gunata, INRA, Montpellier, France. In addition, 3-hydroxy-*α*-ionol, 4-hydroxy-*β*-ionol, 3-hydroxy-*α*-ionone, 4-hydroxy-*β*-ionone, 3-hydroxy-*α*-damascone and 4-hydroxy-*β*-damascone were tested, which were produced by P450-catalyzed biotransformation using *Bacillus megaterium* (Putkaradze et al. 2017). For the substrate screening, 50 µl crude protein extract, 100 mM Tris-HCl buffer (pH 7.5), 600 µM substrate (dissolved in DMSO) and 1 mM UDP-glucose were added to a 200 µl standard assay. The reaction was initiated by the addition of UDP-glucose, incubated with constant shaking at 400 rpm and 30 °C for 17 hours in darkness. The reaction was stopped by heating 10 min at 75 °C. The samples were centrifuged (14,800 rpm) at room temperature for 10 min twice to separate the precipitated protein from the soluble products. Fifty µl of the clear supernatant was analysed by LC-MS and products were monitored using diagnostic ions (Supplemental Table S1). Samples were analyzed with an Agilent 1100 LC/UV system (Agilent Technologies, Waldbronn, Germany) equipped with a reverse-phase column (Luna 3u C18 100A, 150 x 2 mm; Phenomenex) and connected to an Agilent 6340 ion-trap mass spectrometer (Agilent Technologies). A binary gradient consisting of solvent A (water with 0.1% formic acid) and B (methanol with 0.1% formic acid) was performed. The gradient went from 0–3 min 100% A to 50% A; 3–6 min 50% A to 100% B; 6–14 min hold 100% B; 14–14.1 min 100% B to 100% A; 14.1–25 min hold 100% A. The flow rate was 0.2 ml min^-1^ and UV was recorded at 280 nm. MS spectra were acquired in alternating polarity mode and nitrogen was used as nebulizer gas at 30 p.s.i. and as dry gas at 330 °C and 9 L min^-1^. Data were analyzed with Data Analysis 5.1 software (Bruker Daltonics, Bremen, Germany).

### Production of hydroxylated C_13_-apocarotenols

For the large-scale production of hydroxylated norisoprenoids, a CYP109E1 based *B. megaterium* whole-cell system was applied (Putkaradze et al. 2017). C_13_-apocarotenoid (*α*- or *β*-ionol, *α*- or *β*-ionone, *α*- or *β*-damascone) was added to the cell-suspension with a final concentration of 40 mg L^-1^. After 4 h of incubation at 30 °C and 150 rpm, when norisoprenoids were almost completely converted into hydroxyl-products, whole-cell reactions were quenched and extracted with a double volume of ethyl acetate. The organic phase was dried using a rotary evaporator.

### Determination of kinetic constants by UDP Glo™ assay

To determine the enzyme kinetics, the reaction conditions were optimized for each UGT. In order to find the optimal protein amount, 0.5 to 4 μg of UGT709C6, UGT86C10 and UGT73A25; 0.25 to 4 μg of UGT73A24; and 0.5 to 6 μg of UGT72AY1 was examined. The pH optima were determined from pH 4.0 to pH 11.5 using citric acid buffer (50 mM, pH 4.0 to 6.0), sodium phosphate buffer (50 mM, pH 6.0 to 8.0) and Tris-HCl (50 mM, pH 7.5 to 11.5). The optimal temperature was evaluated from 15 to 60 °C in 5 °C-unit intervals. Enzyme kinetics were determined with the UDP-Glo™ assay (Promega, Mannheim, Germany) according to the manufacturer’s instructions using the optimal reaction conditions for each UGT. The reaction mixture (100 μl) contained 50 mM optimal buffer, 100 μM UDP-glucose, a final concentration of 10 to 1200 µM substrate (dissolved in DMSO), and optimal concentration of purified protein. For the blank reaction, DMSO was added instead of the substrate solution. Triplicate analyses were performed. The luminescence was measured with a CLARIOstar Microplate-reader (BMG Labtech, Ortenberg, Germany) and the kinetic constants were calculated by non-linear regression of the obtained enzyme activity employing the Microsoft Excel Solver.

### *α*-Ionyl *β*-D-glucopyranoside production by whole cell biotransformation

Whole cell biotransformation was performed according to the procedure in (Effenberger et al. 2019) using *MpUGT86C10*-pGEX-4T1 for the production of *α*-ionyl *β*-D-glucopyranoside, except that a 5-L-bioreactor was used. Bacteria were grown at 37 °C in M9 minimal medium to an OD_600_ of 1.5. The culture was fed daily with 1% sucrose and 100 µL L^-1^ of *α*-ionol. The yield of *α*-ionyl glucoside in the supernatant at 7 days after IPTG induction was 0.38 g/L. The glucoside produced by the whole-cell biocatalyst could be readily purified by solid phase extraction from the supernatant of the culture. After distillation and ethyl acetate extraction, the colored impurities were removed by activated carbon treatment (Effenberger et al. 2019). LC-MS and NMR analyses confirmed purity and identity, respectively (Supplemental Figure S4).

### NMR spectroscopy of *α*-ionyl glucoside

Thirty mg of pure *α*-ionyl glucoside was dissolved in 600 µl DMSO-D_6_ (99.96%) containing 0.03% (v/v) trimethylsilane (TMS, Sigma-Aldrich, Steinheim, Germany). NMR spectra were recorded with a Bruker MHz Avance III spectrometer (Bruker, Rheinstetten, Germany). A combination of COSY (correlation spectroscopy), HSQC (heteronuclear single quantum coherence), HMBC (heteronuclear multiple-bond correlation), ^1^H, and ^13^C experiments were used for structure elucidation. The ^1^H NMR and ^13^C NMR spectra were recorded at 400.133 MHz and 100.624 MHz, respectively. The chemical shifts were referred to the solvent signal and TMS. The spectra were acquired and processed with MestReNova software (https://mestrelab.com/).

### Agroinfiltration of UGTs into *N. benthamiana* leaves

Agroinfiltration was performed according to (Sun et al. 2019) using a viral vector system to deliver the various modules into plant cells. The full-length open reading frames (ORF) of *UGT709C6, UGT86C10, UGT72AY1, UGT73A24* and *UGT73A25* were amplified by PCR from plasmid pGEX-4T1-UGTs using the primers listed in Supplemental Table S4. The PCR products were double digested with restriction enzymes (Supplemental Table S4), and ligated into pICH11599 vectors, according to (Sun et al. 2019) to yield pICH11599-UGTs. The recombinant genes were subjected to sequencing to confirm the sequence of the inserts. *Agrobacterium* strains carrying pICH17388, pICH14011 and pICH11599-UGTs were mixed and infiltrated into *N. benthamiana* leaves. An empty pICH11599 vector was infiltrated and served as negative control. The infiltrated plants grew under the same conditions. Leaves were sampled 7 and 10 days after infiltration. Proteins were extracted from infiltrated leaves and analyzed for enzyme activity by LC-MS according to (Sun et al. 2019).

### Verification of *UGT* overexpression by quantitative real-time PCR analysis

Quantitative real-time PCR (qRT-PCR) analysis was performed on *N. benthamiana* leaves 7 d (NbUGTs) and 10 d (MpUGTs) after agroinfiltration according to (Sun et al. 2019). Actin was used as internal control for normalization (Supplemental Table S4). The reaction was run on a StepOnePlus™ system (Applied Biosystems™ Waltham, MA, USA). Subsequently, 2% agarose gel electrophoresis was applied to confirm that the desired amplicons had been generated, and the relative expression level was analyzed by applying a modified 2 ^ΔΔ-Ct^ method taking reference genes and gene specific amplification efficiencies into account (Sun et al. 2019).

### Aglycone library generation

To identify natural apocarotenoid substrates of the UGTs, aglycone libraries were prepared from tobacco leaves as described by (Sun et al. 2019) and from *M.× piperita* leaves. One hundred and eighty g mint leaves were homogenized in 500 ml water, subjected to XAD2 (237 g) solid phase extraction and glycosides were eluted with 50 ml methanol. Methanol was removed by rotary evaporation, the residue dissolved in 10 ml water, and 10 ml citrate/phosphate buffer (pH 4.3) containing 1.6 g of the glycosidase Rapidase 2000 (DSM, Düsseldorf, Germany) was added. Hydrolysis was performed at room temperature, overnight. Aglycones were extracted with 20 ml ethyl acetate, concentrated to 1 ml (aglycone library) and analyzed by GC-MS according to (Huang et al. 2009; Schmidt et al. 2006). The library also served as substrate in enzyme assays with MpUGT86C10 and an empty vector control. To avoid loss of volatiles extracts were carefully concentrated by using a Vigreux column.

### Co-infiltration of Agrobacterium and acceptor substrates

*N.benthamiana* (8 weeks old) leaves were agroinfiltrated with *MpUGT86C10* and pICH11599 (empty vector), respectively as described before. The model apocarotenoid substrates *α*-ionol and *β*-ionol were dissolved in DMSO (8.3%) and dropped (100 µL in total) adaxial onto *N. benthamiana* leaves using a pipette within 5 min, 3 days after agroinfiltration. Wild type leaves and leaves agroinfiltrated with an empty vector control were also treated with the same apocarotenoid substrates. Leaves were collected 5 and 7 days after substrate infiltration. Proteins were extracted from infiltrated leaves for the analysis of enzyme activity and metabolites were extracted for the identification of glucoside products by LC-MS.

### Germination test using *N. benthamiana* seeds

*N. benthamiana* seeds were washed with 75% ethanol, 3.5% NaClO and H_2_O, before being placed on a wetted filter paper to germinate at room temperature and a 16 h light, 8 h dark photoperiod. To determine the effect of C_13_-apocarotenoids on seed germination, a 1.5 ml solution of *α*-ionol and *α*-ionyl glucoside (0.1; 1; 10 and 100 mM, respectively) in 1% DMSO were tested. Twenty seeds per treatment were used. Experiment was conducted in triplicate. After two weeks of incubation time, the germination rates and embryo sizes were measured.

### Accession numbers

NbUGT72AY1, Nbv6.1trP2283; NbUGT73A24, Nb6.1trP32845; NbUGT73A25, Nb6.1trP48287; NbUGT85A73, Nb6.1trP20189.

## Acknowledgements

We are thankful to Michael Strebl, Maximilian Merz, and Isabelle Effenberger for their technical support. Thanks to Alexander Christmann for providing tobacco plants. The authors declare no competing interests.

## Funding

We thank the China Scholarship Council (CSC) (no. 201506060185), International Graduate School of Science and Engineering (IGSSE) (no. 10.05) and Deutsche Forschungsgemeinschaft DFG SCHW 634/32-1 for funding.

The data that support the findings of this study are available from the corresponding author upon reasonable request.

Short legends for Supplemental Figures and Tables

**Figure S1.** Chemical structures of selected C_13_-apocarotenoids (norisoprenoids) isolated from plants.

**Figure S2.** Relative expression levels of *UGTs* in different tissues of *N. benthamiana* and *M. x piperita*.

Figure S3. SDS-PAGE analysis of six UGTs.

**Figure S4.** Heteronuclear Single Quantum Correlation (HSQC) Nuclear Magnetic Resonance (NMR) spectroscopy of enzymatically synthesized *α*-ionyl *β*-D-glucopyranoside.

**Figure S5.** Phylogenetic tree of the analyzed UGTs from *N. benthamiana* and *M. x piperita*.

Figure S6. Amino acid sequence alignment of selected UGTs from *M. x piperita and N. benthamiana* that glucosylated C_13_-apocarotenols.

**Figure S7.** GC-MS of apocarotenoids formed by P450-mediated biotransformation of α-ionol to produce 3-hydroxy-α-ionol and LC-MS analysis to detect 3-hydroxy-α-ionyl glucoside produced by UGTs.

**Figure S8.** GC-MS of apocarotenoids formed by P450-mediated biotransformation of β-ionol to produce 4-hydroxy-β-ionol and LC-MS analysis to detect 4-hydroxy-β-ionyl glucoside produced by UGTs.

**Figure S9.** GC-MS of apocarotenoids formed by P450-mediated biotransformation of α-ionone to produce 3-hydroxy-α-ionone and LC-MS analysis to detect 3-hydroxy-α-ionone glucoside produced by UGTs.

**Figure S10.** GC-MS of apocarotenoids formed by P450-mediated biotransformation of β-ionone to produce 4-hydroxy-β-ionone and LC-MS analysis to detect 4-hydroxy-β-ionone glucoside produced by UGTs.

**Figure S11.** GC-MS of apocarotenoids formed by P450-mediated biotransformation of α-damascone to produce 3-hydroxy-α-damascone and LC-MS analysis to detect 3-hydroxy-α-damascone glucoside produced by UGTs.

**Figure S12.** GC-MS of apocarotenoids formed by P450-mediated biotransformation of β-damascone to produce 4-hydroxy-β-damascone and LC-MS analysis to detect 4-hydroxy-β-damascene glucoside produced by UGTs.

**Figure S13.** Reconstitution of apocarotenoid glucosides by MpUGT86C10 from a mint aglcone library.

**Figure S14.** Fraction 10 of the tobacco aglycone library was used as substrate source for recombinant NbUGT72AY1 from *N. benthamiana.*

**Figure S15.** LC-MS analysis (combined extracted ion chromatograms *m/z* -401 and - 391) of products (ionyl glucosides) obtained by enzyme activity assays with protein extracts from wild type plants.

**Figure S16.** LC-MS analysis (combined extracted ion chromatograms *m/z* -401 and - 391) of products (ionyl glucosides) obtained by enzyme activity assays with protein extracts from control plants (agroinfiltrated with an empty vector).

**Figure S17.** LC-MS analysis (combined extracted ion chromatograms *m/z* -401 and - 391) of products (ionyl glucosides) obtained by enzyme activity assays with protein extracts from *MpUGT86C10*–infiltrated plants.

**Table S1.** Apocarotenoids used in the study, their molecular weights (MW), as well as the molecular weights and diagnostic ions of their glucosides.

**Table S2.** UGTs amino acids sequence identities (%).

**Table S3.** The optimal reaction conditions for each UGT for the kinetic assay determined with UDP Glo™ assay.

**Table S4.** Primers used for amplification of the UGT genes and overexpression qRT-PCR. fwd = forward, rev = reverse

