## Supplemental Figures and Tables for "Glycosylation of bioactive C_13_-apocarotenols in *Nicotiana benthamiana* and *Mentha × piperita*": Supporting Information Tables and Figures 2020.pdf

### Figures S1 - S17
### Tables S1 - S4

### FIGURES

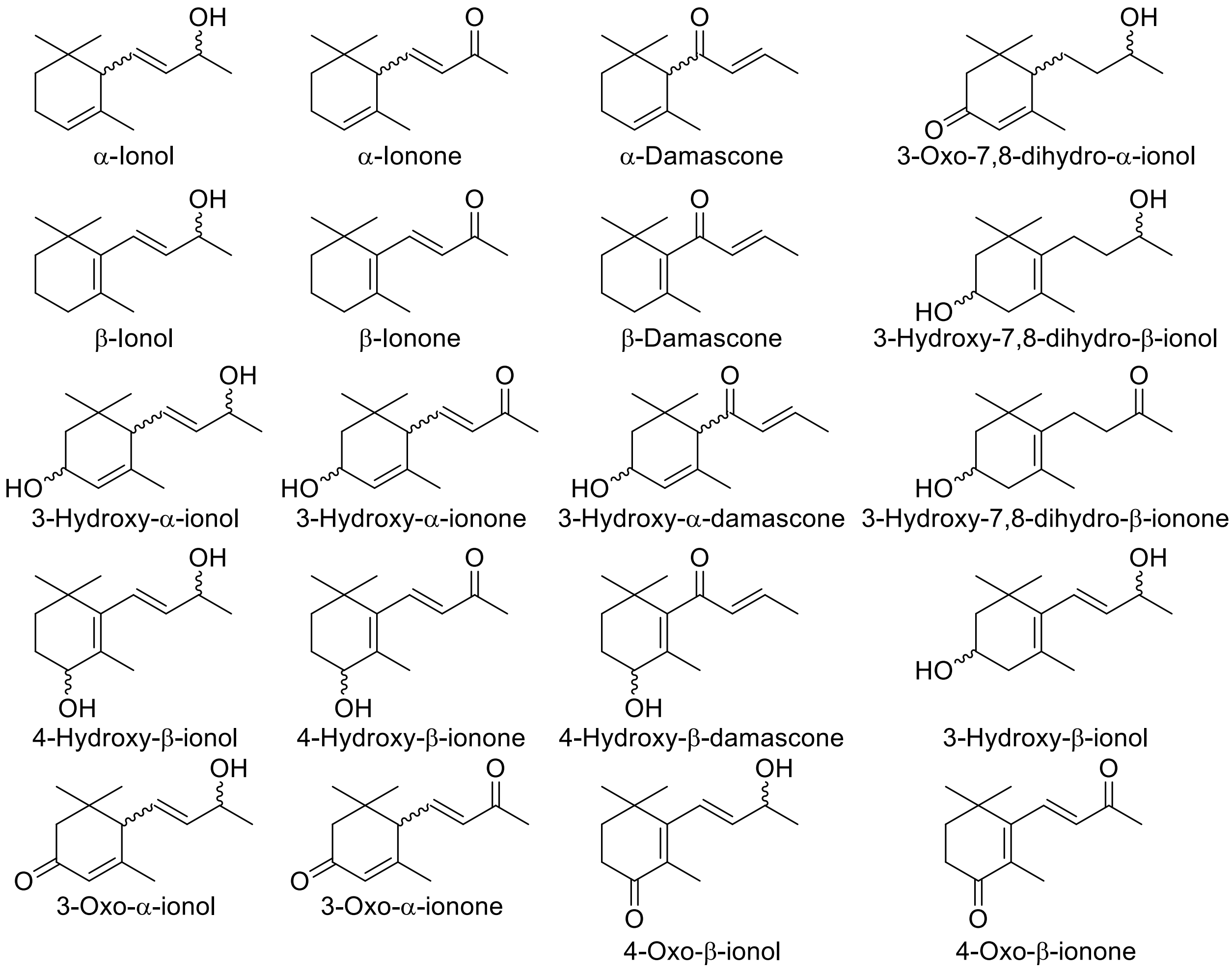


**Figure S1. Chemical structures of selected $C_{13}$-apocarotenoids (norisoprenoids) isolated from plants**.

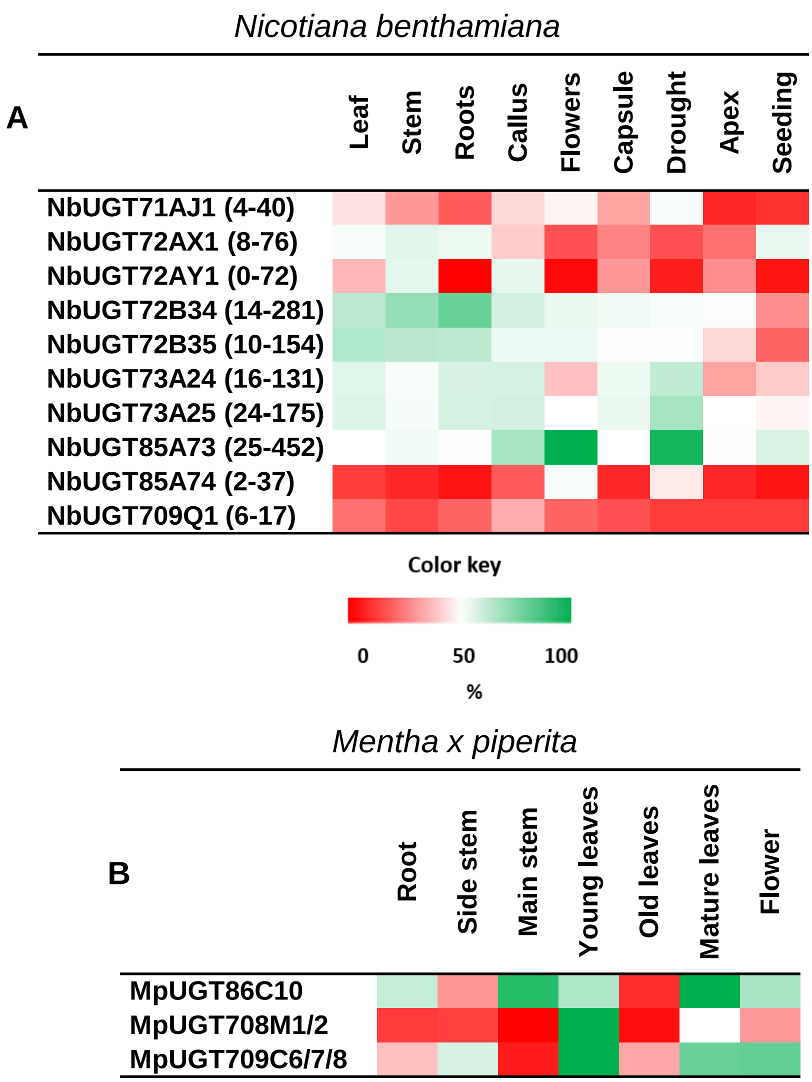


**Figure S2. Relative expression levels of *UGTs* in different tissues of *N. benthamiana* and *M. x piperita*.** UGT expression data in percent max [%]. Heat maps show relative expression in *N. benthamiana* and *M. x piperita* tissues with lowest levels in red and highest levels in green. (A) Individual min. and max. transcript numbers in reads/million are given in parentheses. Data from: Nakasugi K, Crowhurst R, Bally J, Waterhouse P (2014) Combining Transcriptome Assemblies from Multiple De Novo Assemblers in the Allo-Tetraploid Plant *Nicotiana benthamiana*. PLoS ONE 9(3): e91776. doi:10.1371/journal.pone.0091776 and http://www.benthgenome.com (B) Expression levels of UGTs determined by qRT-PCR analysis. *Actin* (KM044035.1) served as reference gene and the lowest and highest transcript level of a gene in a tissue was set at 0 and 100%, respectively. Leaves of 0.5 to 1.5 cm length were harvested as young leaves. The 5th leaf pair from the top constituted mature leaves and leaves below (7$^{th}$ pair) were considered as old leaves.

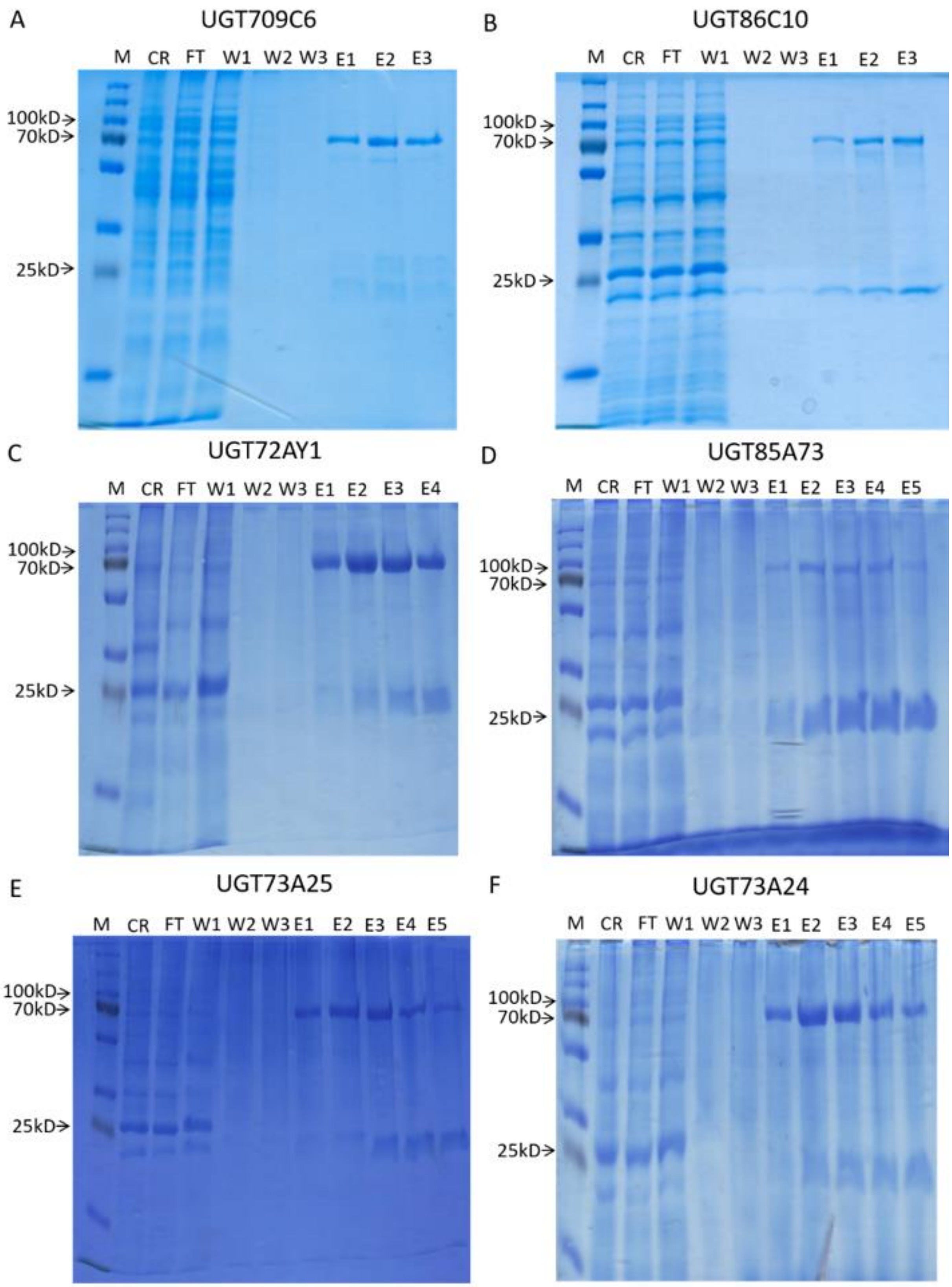


**Figure S3. SDS-PAGE analysis of six selected UGTs.** UGT709C6 (A), UGT86C10 (B), UGT72AY1 (C), UGT85A73 (D), UGT73A25 (E), and UGT73A24 (F). M molecular marker, CR crude protein extract; FT low-through; W1–3 wash 1–3; E1–5 elution 1–5. For further information refer to Sun *et al*., 2019.

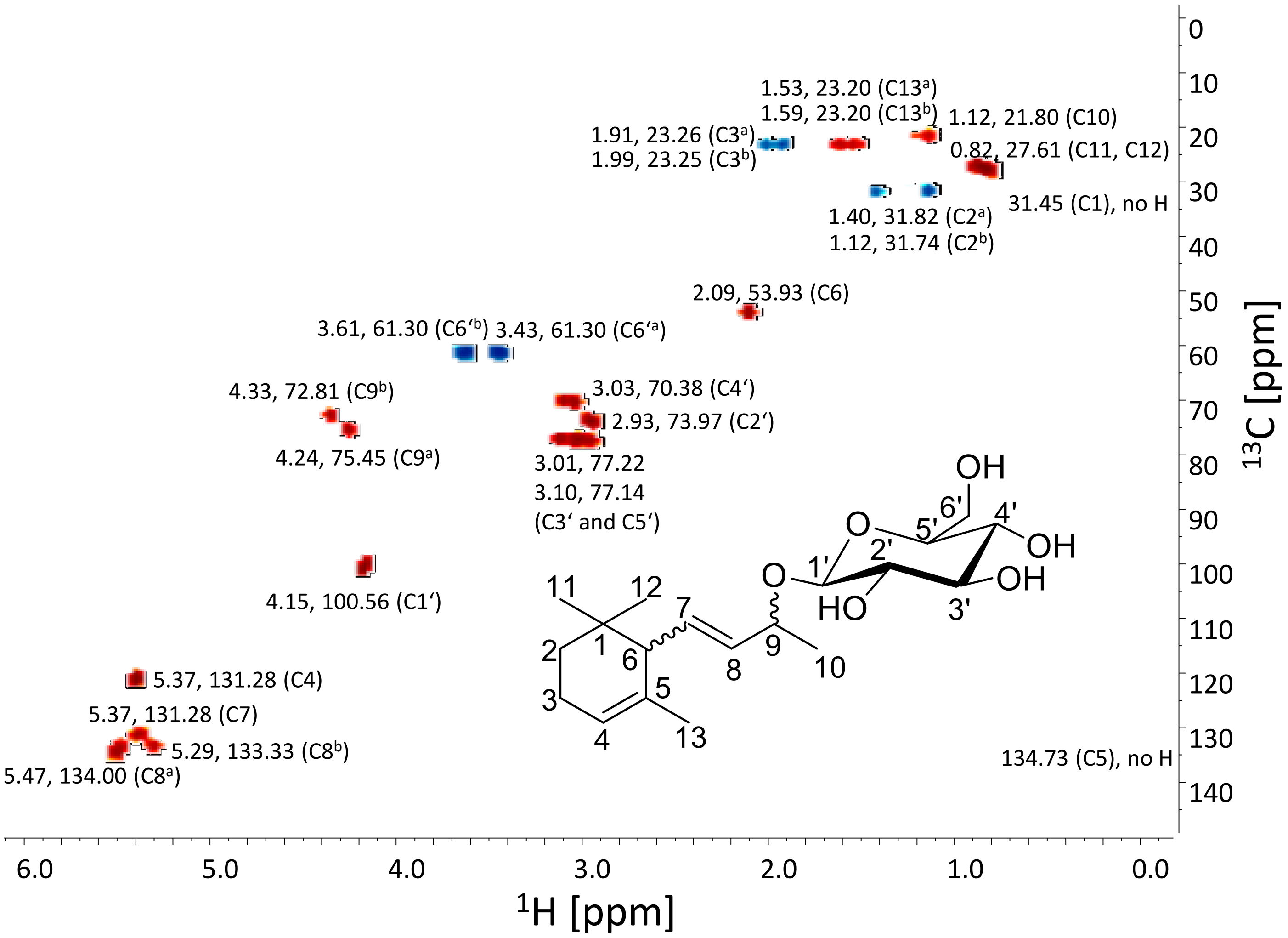


**Figure S4. Heteronuclear Single Quantum Correlation (HSQC) Nuclear Magnetic Resonance (NMR) spectroscopy of enzymatically synthesized *α*-ionyl *β*-D-glucopyranoside.** The sample was evaporated and dissolved in 600 µl DMSO-D6 (99.9%) containing 0.1% (v/v) trimethylsilane (TMS). NMR spectra were recorded with a Bruker DRX 400 spectrometer (Bruker, Karlsruhe, Germany). The chemical shifts were referred to the solvent signal and TMS. The spectra were acquired and processed with MestReNova software.

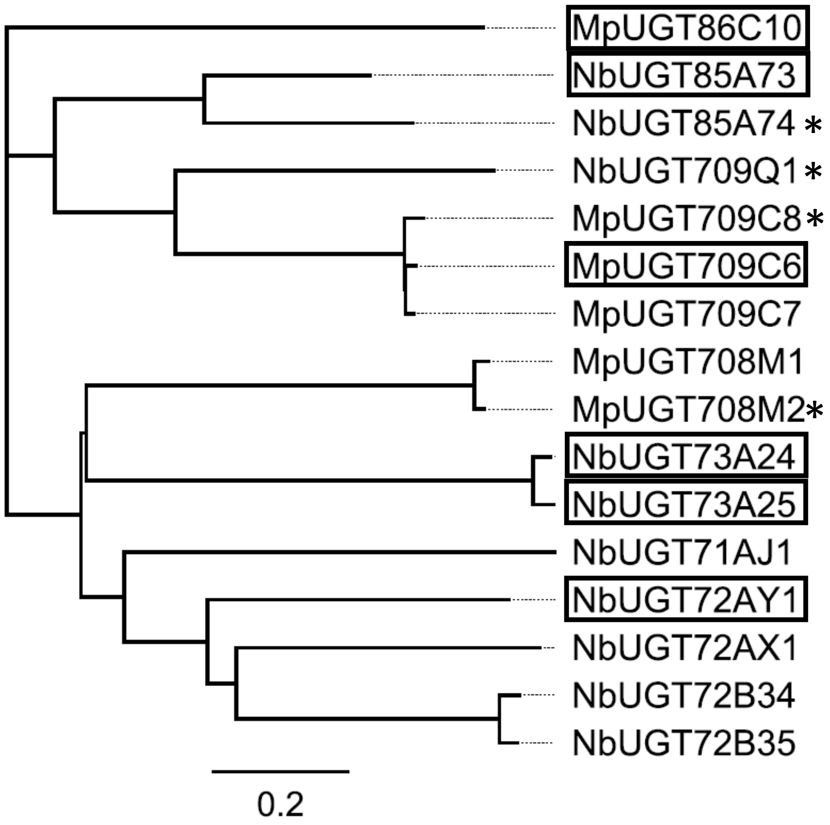


**Figure S5. Phylogenetic tree of the analyzed UGTs from *N. benthamiana* and *M. x piperita*.** The tree was constructed with Geneious using default values (Genetic distance model Jukes-Cantor; Neighbor-Joining tree build method, no outgroup; bootstrap 1,000 trials). Enzymes catalyzing the glucosylation of apocarotenoids are boxed.* UGTs are most probably inactive as important features of UGTs are missing.

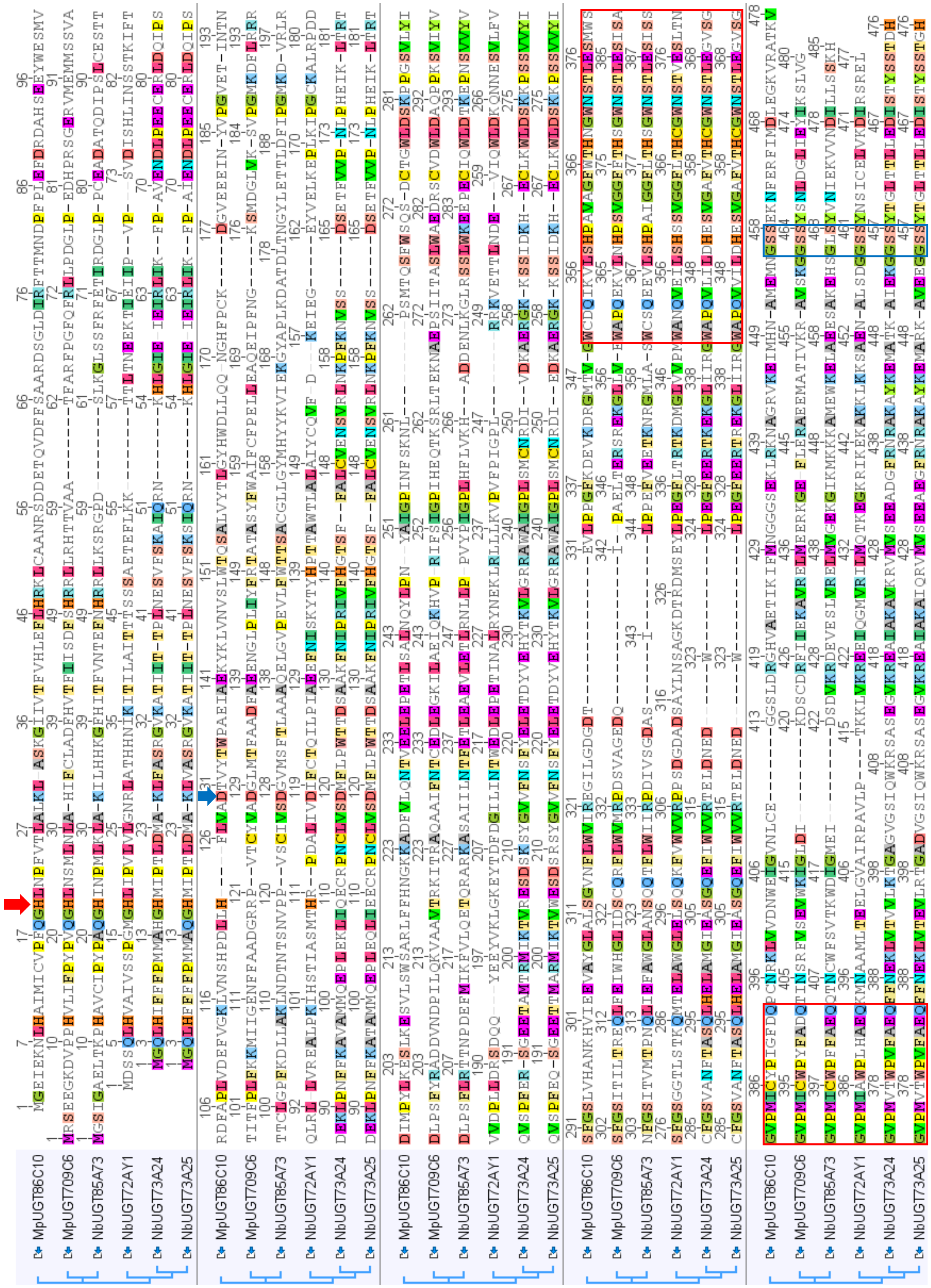


**Figure S6. Amino acid sequence alignment of selected UGTs from *M. x piperita and N. benthamiana* that glucosylated $C_{13}$-apocarotenols.** Red and blue arrows show the catalytically active His and Asp, respectively and the red and blue box the conserved PSPG box and GSS motif, respectively. Default values of the Geneious Pro 5.5.6 program (http://www.geneious.com/) were used.

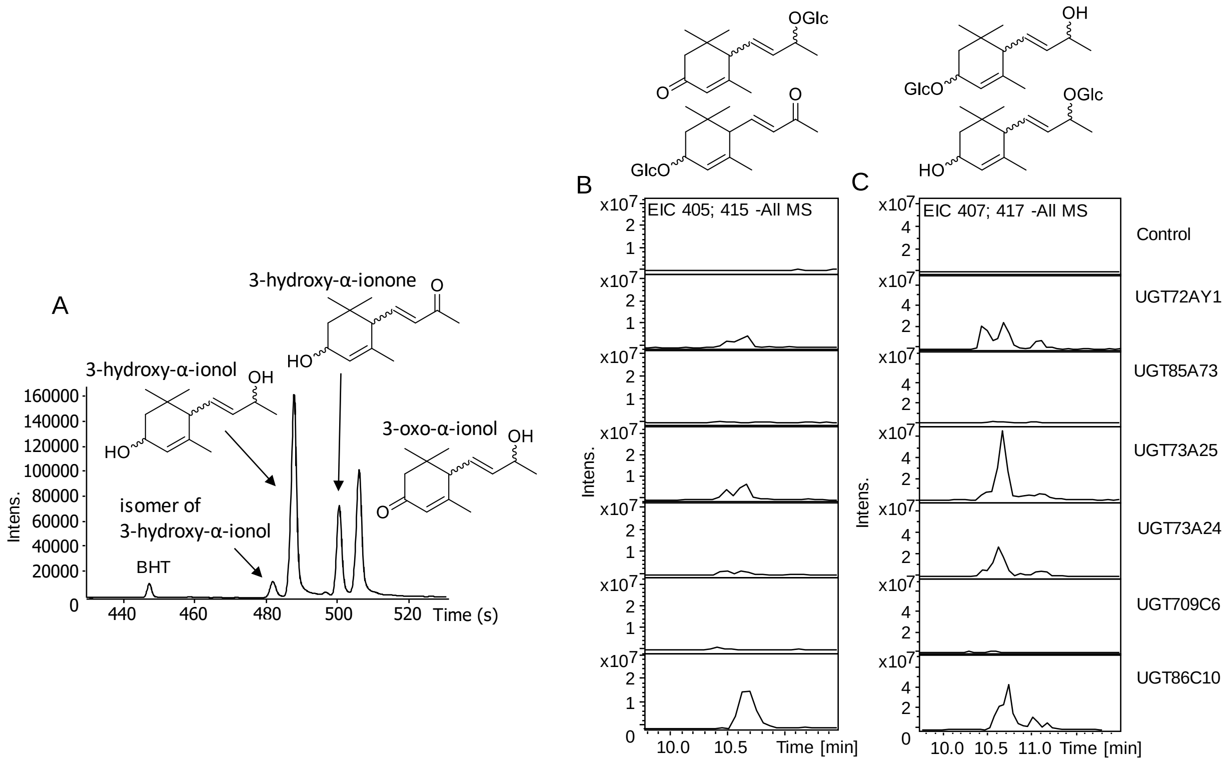


**Figure S7. GC-MS of apocarotenoids formed by P450-mediated biotransformation of α-ionol to produce 3-hydroxy-α-ionol and LC-MS analysis to detect 3-hydroxy-α-ionyl glucoside produced by UGTs.** The GC-MS analysis (A) reveals three apocarotenoids, BHT butylated hydroxytoluene. The LC-MS (B) extracted ion chromatograms (EIC) at *m/z* 405 and 415 show 3-hydroxy-α-ionone glucoside and 3-oxo-α-ionyl glucoside while *m/z* 407 and 417 show isomeric 3-hydroxy-α-ionyl glucosides.

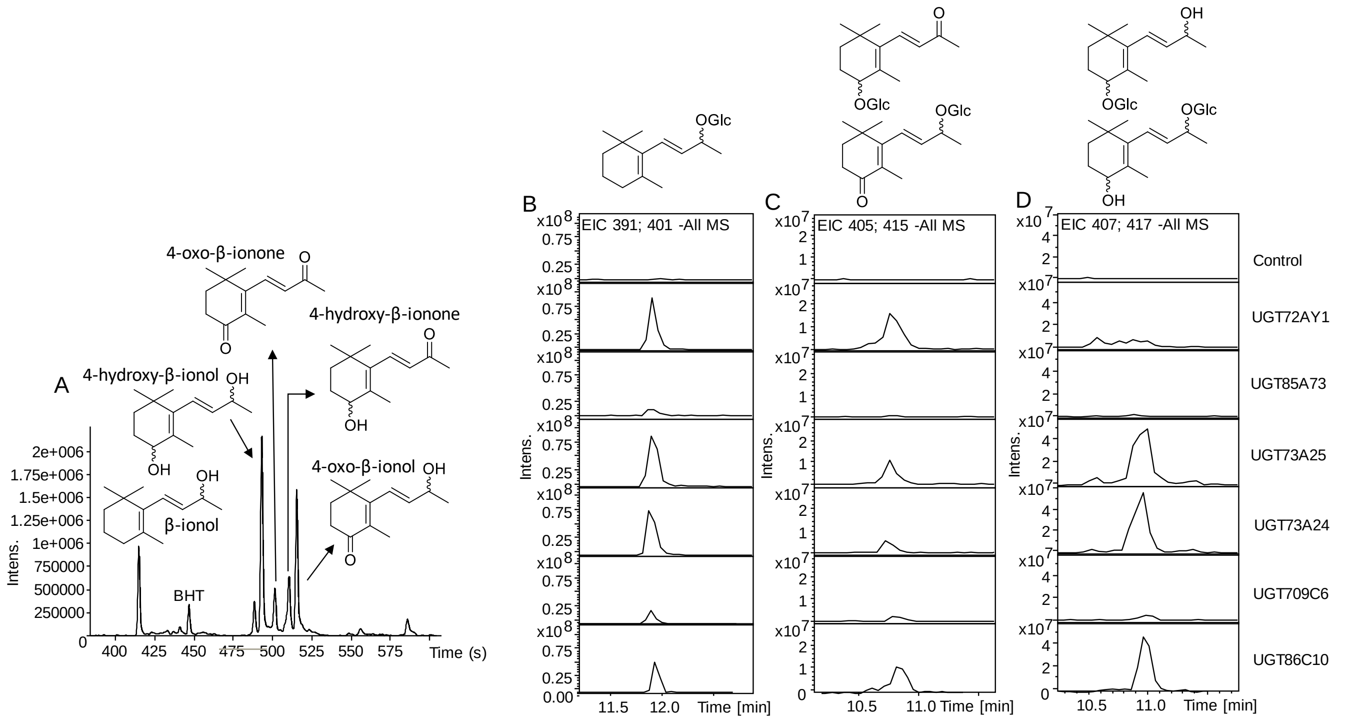


**Figure S8. GC-MS of apocarotenoids formed by P450-mediated biotransformation of β-ionol to produce 4-hydroxy-β-ionol and LC-MS analysis to detect 4-hydroxy-β-ionyl glucoside produced by UGTs.** The GC-MS analysis (A) reveals five apocarotenoids, BHT butylated hydroxytoluene. The LC-MS (B) extracted ion chromatograms (EIC) at *m/z* 391 and 401 show β-ionyl glucoside, *m/z* 405 and 415 show 4-oxo-β-ionyl glucoside and 4-hydroxy-β-ionone glucoside while *m/z* 407 and 417 show isomeric 4-hydroxy-β-ionyl glucosides.

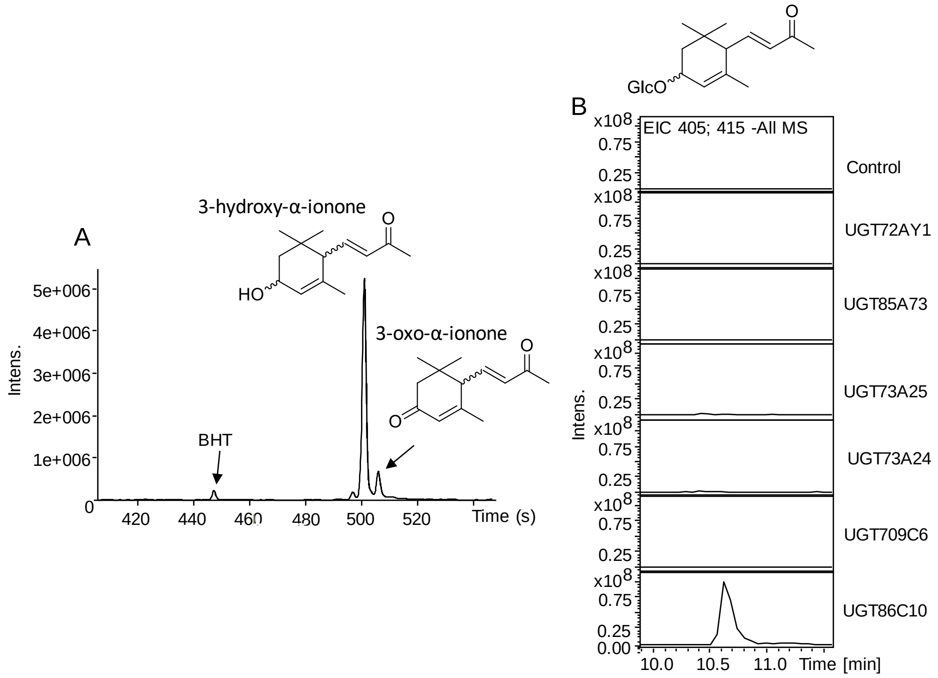


**Figure S9. GC-MS of apocarotenoids formed by P450-mediated biotransformation of α-ionone to produce 3-hydroxy-α-ionone and LC-MS analysis to detect 3-hydroxy-α-ionone glucoside produced by UGTs.** The GC-MS analysis (A) reveals two apocarotenoids, BHT butylated hydroxytoluene. The LC-MS (B) extracted ion chromatograms (EIC) at *m/z* 405 and 415 show 3-hydroxy-α-ionone glucoside.

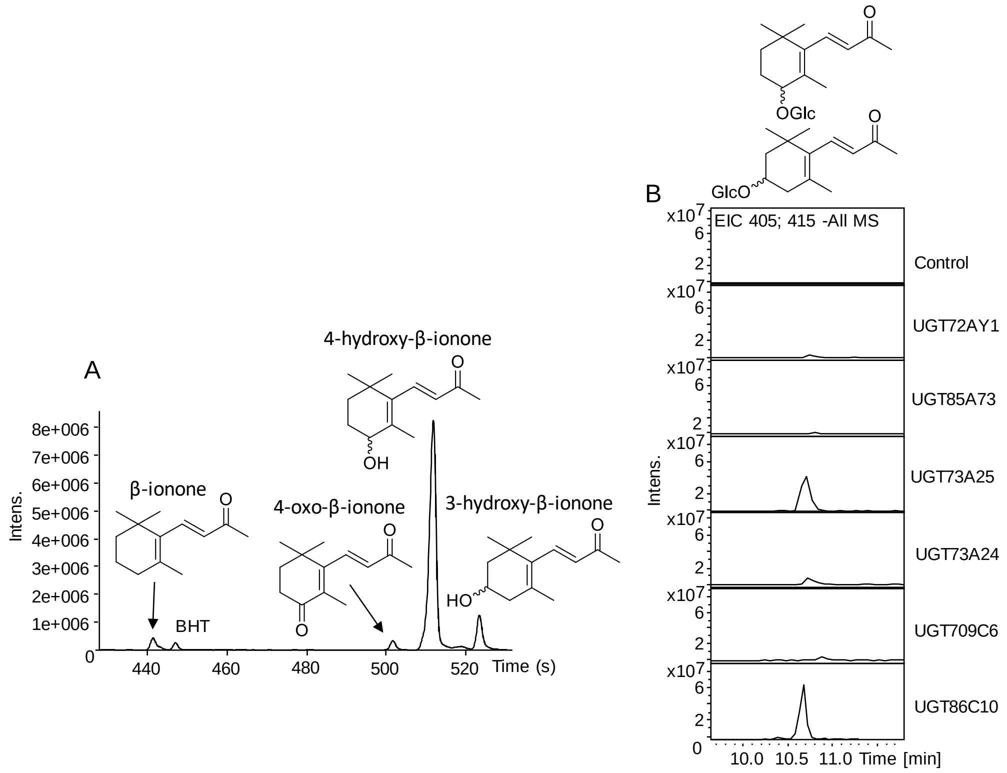


**Figure S10. GC-MS of apocarotenoids formed by P450-mediated biotransformation of β-ionone to produce 4-hydroxy-β-ionone and LC-MS analysis to detect 4-hydroxy-β-ionone glucoside produced by UGTs.** The GC-MS analysis (A) reveals four apocarotenoids, BHT butylated hydroxytoluene. The LC-MS (B) extracted ion chromatograms (EIC) at *m/z* 405 and 415 show 4-hydroxy-β-ionyl glucoside and 3-hydroxy-β-ionyl glucoside.

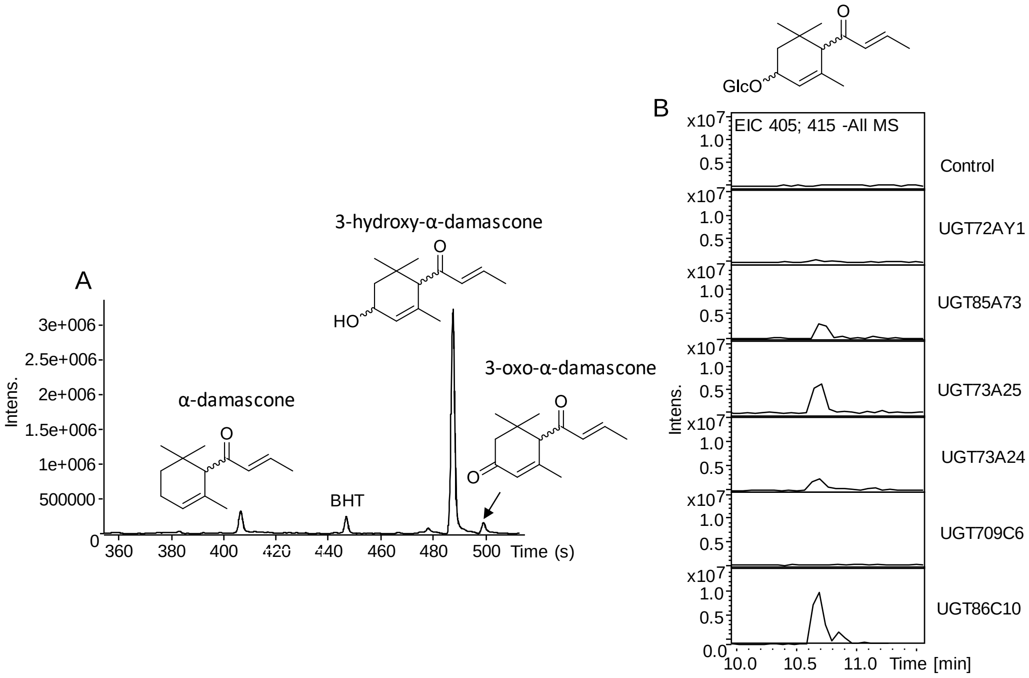


**Figure S11. GC-MS of apocarotenoids formed by P450-mediated biotransformation of α-damascone to produce 3-hydroxy-α-damascone and LC-MS analysis to detect 3-hydroxy-α-damascone glucoside produced by UGTs.** The GC-MS analysis (A) reveals three apocarotenoids, BHT butylated hydroxytoluene. The LC-MS (B) extracted ion chromatograms (EIC) at *m/z* 405 and 415 show 3-hydroxy-α-damascone glucoside.

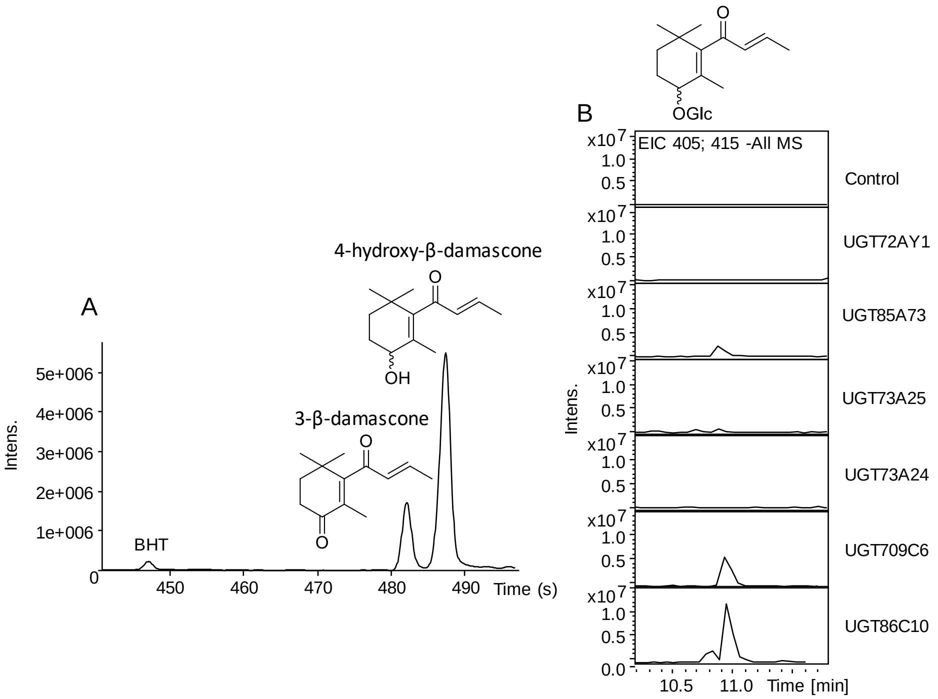


**Figure S12. GC-MS of apocarotenoids formed by P450-mediated biotransformation of β-damascone to produce 4-hydroxy-β-damascone and LC-MS analysis to detect 4-hydroxy-β-damascene glucoside produced by UGTs.** The GC-MS analysis (A) reveals two apocarotenoids, BHT butylated hydroxytoluene. The LC-MS (B) extracted ion chromatograms (EIC) at *m/z* 405 and 415 show 4-hydroxy-β-damascone glucoside.

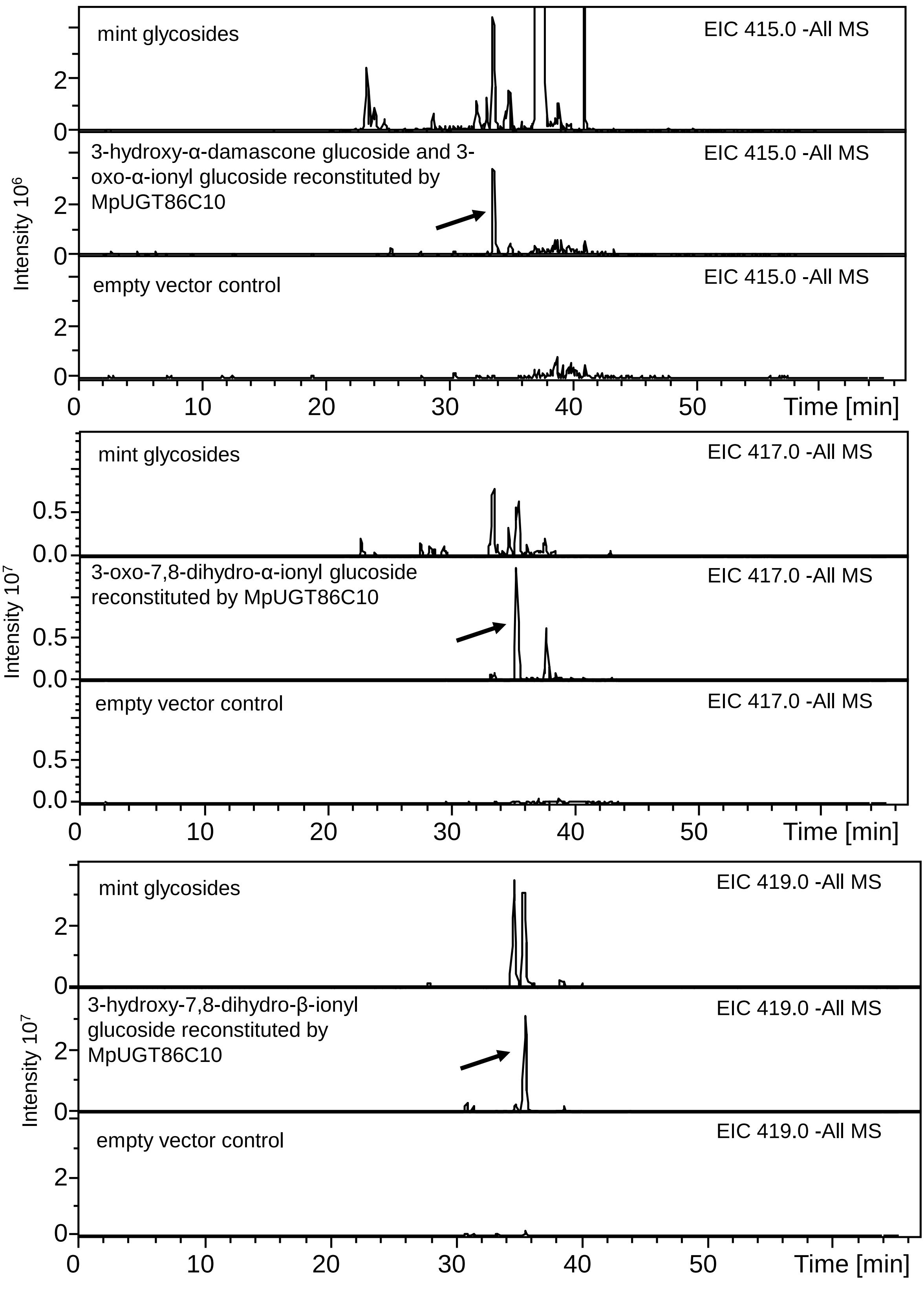


**Figure S13. Reconstitution of apocarotenoid glucosides by MpUGT86C10 from a mint aglcone library.** A glycoside extract from *M. x piperita* leaves served as reference (top panel), reconstituted glycosides by MpUGT86C10 (middle panel) and empty vector control (lower level).

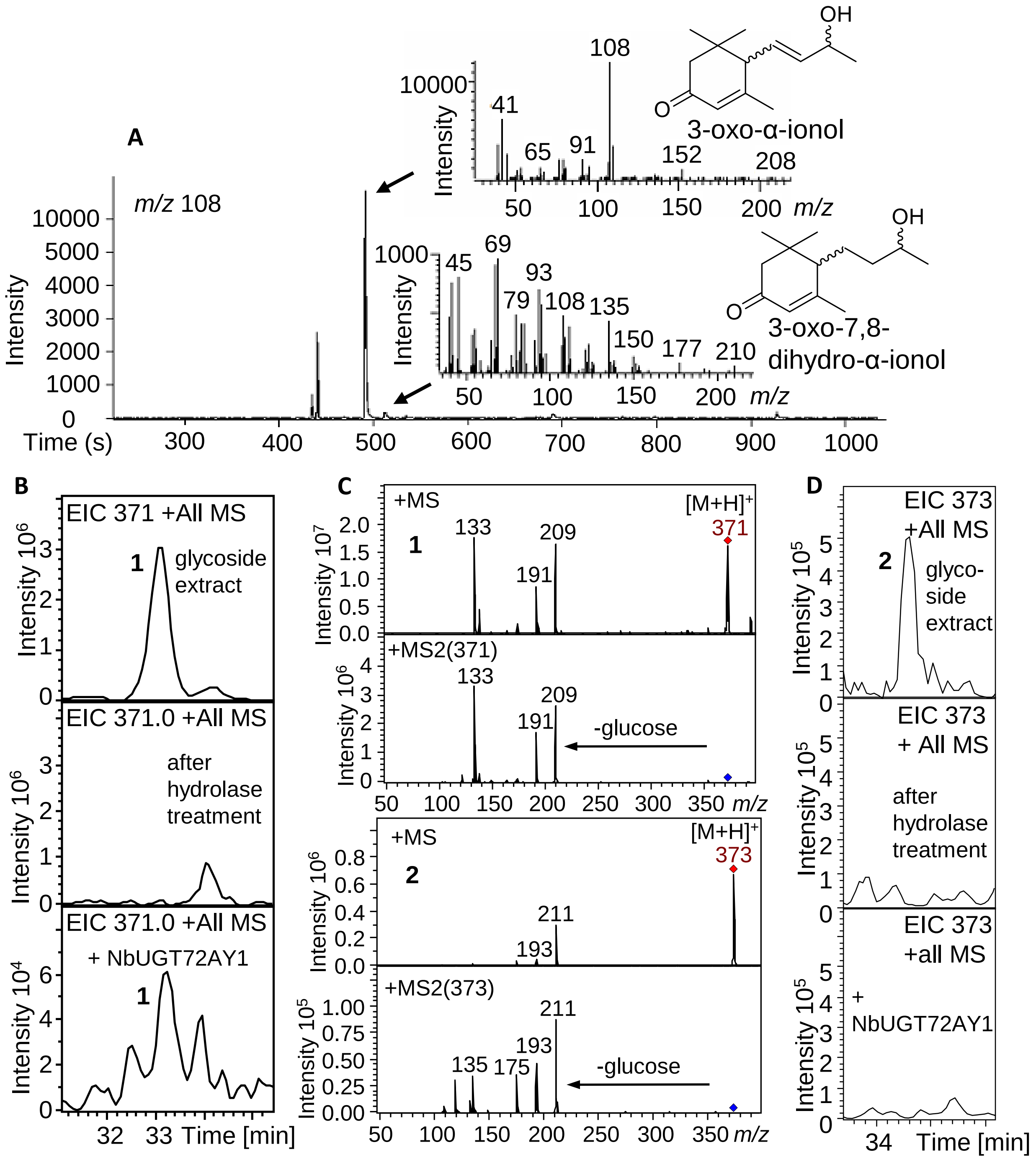


**Figure S14. Fraction 10 of the tobacco aglycone library was used as substrate source for recombinant NbUGT72AY1 from *N. benthamiana*.** 3-Oxo-α-ionol and 3-oxo-7,8-dihydro-α-ionol were released by glucosidase (Rapidase) from the tobacco glycoside extract (aglycone library) and identified by GC-MS (A). Glycosides of fraction 10 of the tobacco extract were analyzed by LC-MS (B, D) and 3-oxo-α-ionyl- (1) and 3-oxo-7,8-dihydro-α-ionyl glucoside (2) were putative identified by their mass spectra (C). Both glucosides were not observed after hydrolase treatment (B, D middle panel) and 1 was reconstituted by NbUGT72AY1 (B, lower panel) while 2 was not reconstituted (D, lower panel).

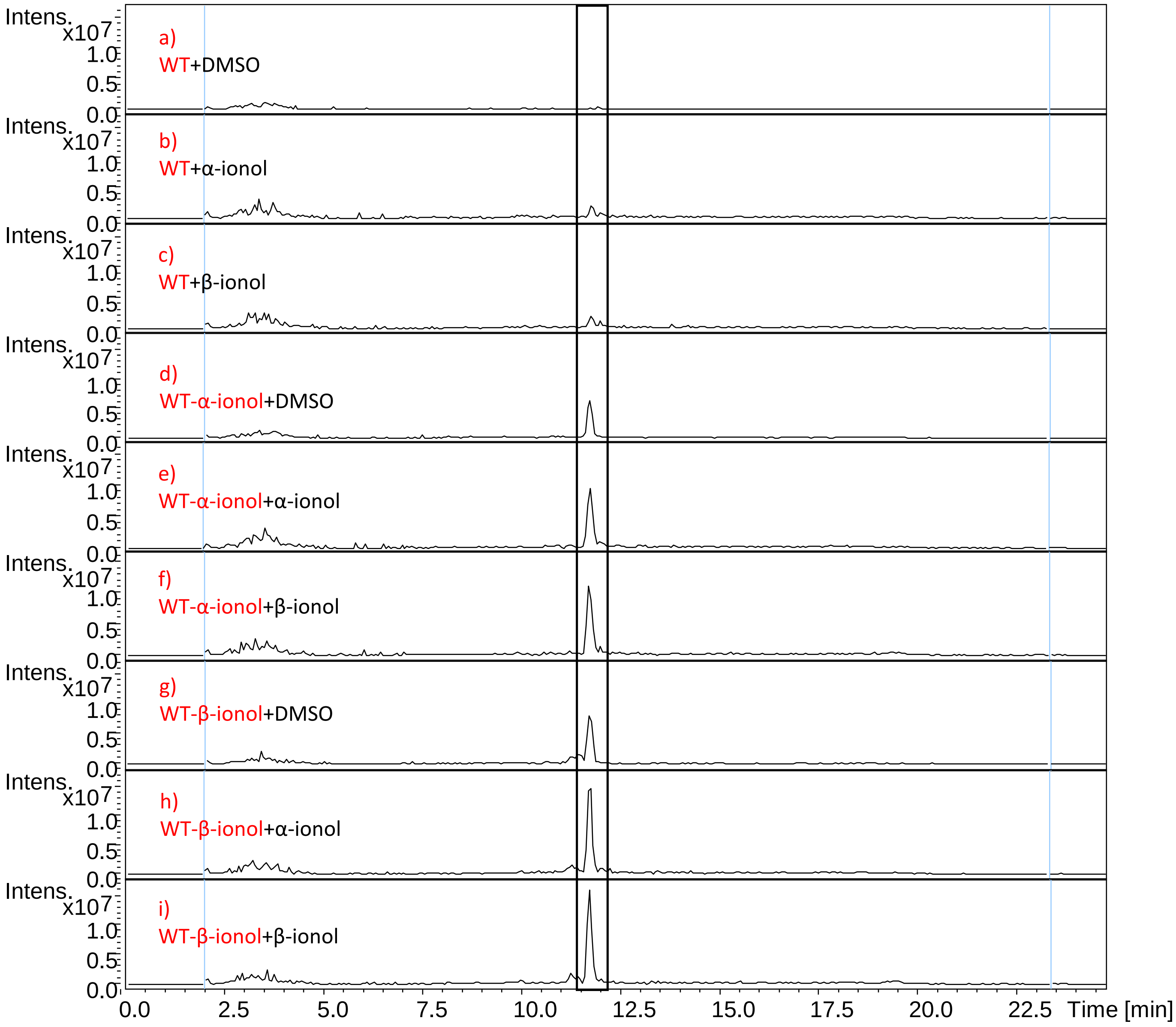


**Figure S15. LC-MS analysis (combined extracted ion chromatograms *m/z* -401 and -391) of products (ionyl glucosides) obtained by enzyme activity assays with protein extracts from wild type plants.** Protein extracts were incubated for 2 hr at 30 °C with the solvent dimethyl sulfoxide (DMSO, as control), α-ionol and β-ionol (a-c). Protein extracts were also isolated from WT leaves, which were pretreated for 5 days with DMSO, α-ionol and β-ionol while still attached to the plants. The protein extracts isolated from these leaves were also incubated with DMSO, α-ionol and β-ionol (d-i).

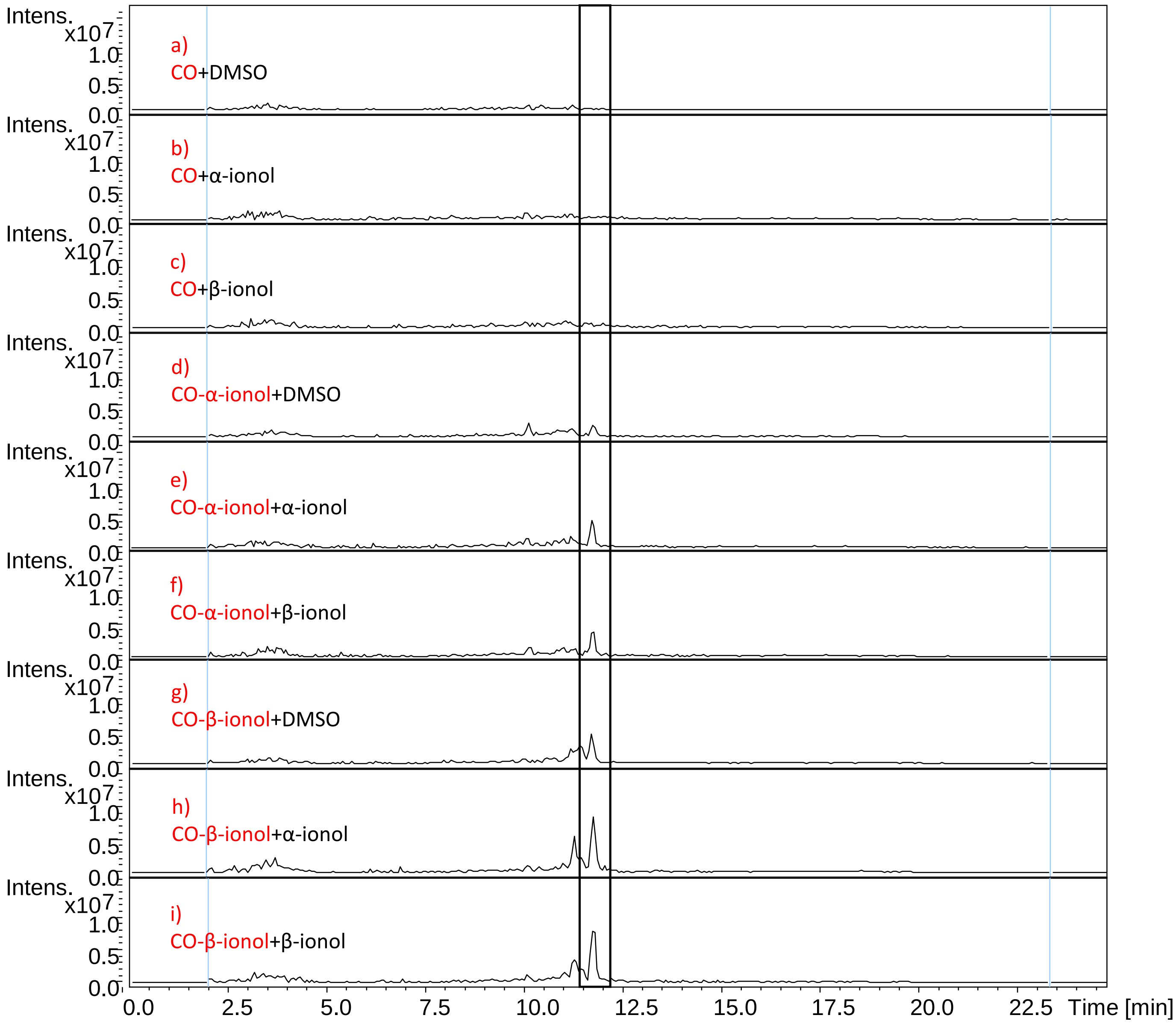


**Figure S16. LC-MS analysis (combined extracted ion chromatograms *m/z* -401 and -391) of products (ionyl glucosides) obtained by enzyme activity assays with protein extracts from control plants (agroinfiltrated with an empty vector).** Protein extracts were incubated for 2 hr at 30 °C with the solvent dimethyl sulfoxide (DMSO, as control), α-ionol and β-ionol (a-c). Protein extracts were also isolated from CO leaves, which were pretreated for 5 days with DMSO, α-ionol and β-ionol while still attached to the plants. The protein extracts isolated from these leaves were also incubated with DMSO, α-ionol and β-ionol (d-i).

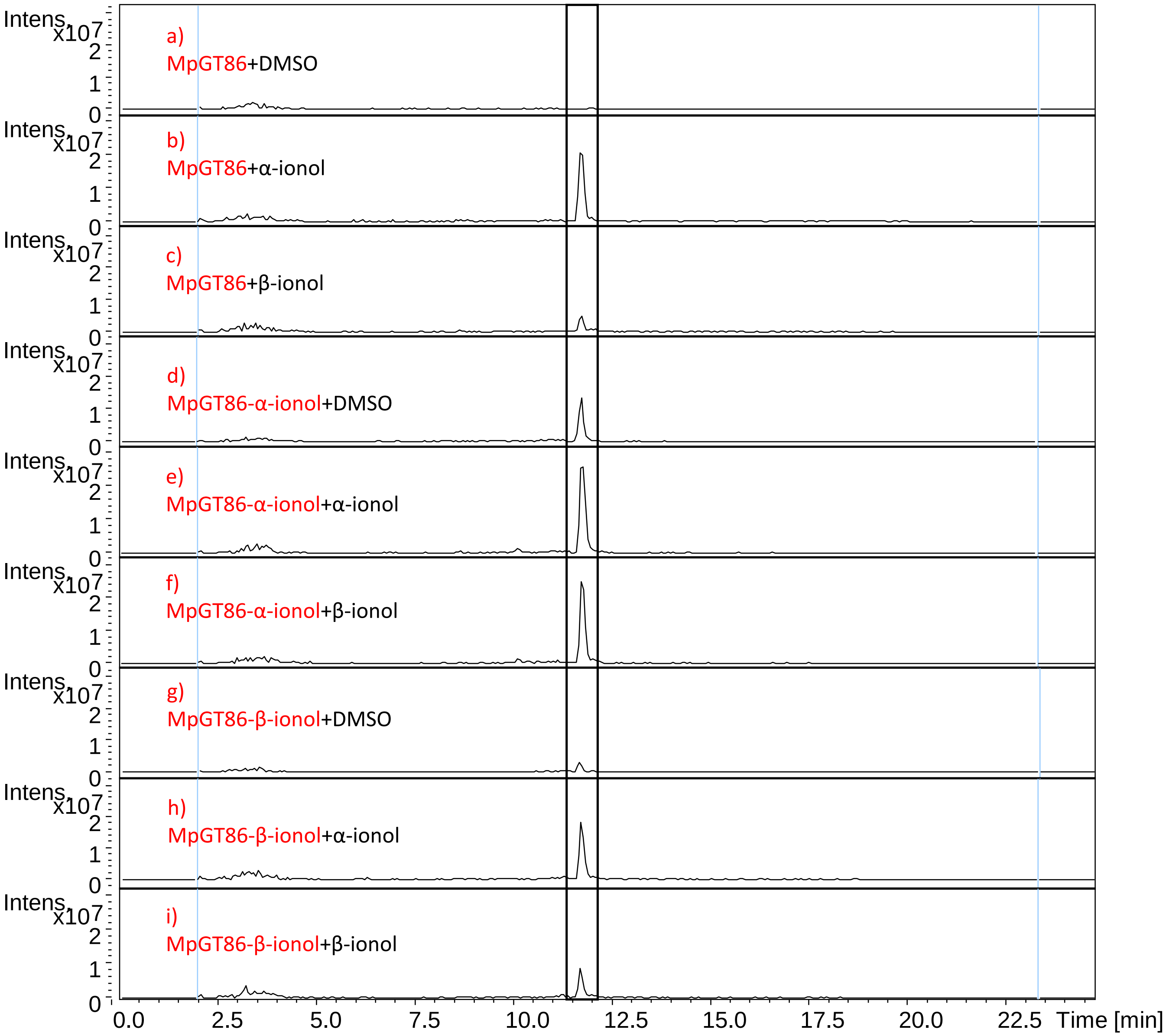


**Figure S17. LC-MS analysis (combined extracted ion chromatograms *m/z* -401 and -391) of products (ionyl glucosides) obtained by enzyme activity assays with protein extracts from *MpUGT86C10*–infiltrated plants.** Protein extracts were incubated for 2 hr at 30 °C with the solvent dimethyl sulfoxide (DMSO, as control), α-ionol and β-ionol (a-c). Protein extracts were also isolated from *MpUGT86C10*–infiltrated leaves, which were pretreated for 5 days with DMSO, α-ionol and β-ionol while still attached to the plants. The protein extracts isolated from these leaves were also incubated with DMSO, α-ionol and β-ionol (d-i).

### TABLES

**Table S1.** Apocarotenoids used in the study, their molecular weights (MW), as well as the molecular weights and diagnostic ions of their glucosides.

| | | | diagnostic ions of the glucoside | | | | | |
|---|---|---|---|---|---|---|---|---|
| | MW aglycone | MW glucoside | [ M+H]+ | [ M+Na]+ | [M-H]- | [ M+Cl]- | [M+HCOO]- | [2M-H]- |
| α-Ionol/β-ionol | 194 | 356 | 357 | 379 | 355 | 391 | 401 | 711 |
| 7,8-dihydro-α-ionol/ 7,8-dihydro-β-ionol | 196 | 358 | 359 | 381 | 357 | 393 | 403 | 713 |
| 3-Hydroxy-α-ionone/ 4-hydroxy-β-ionone/ 3-hydroxy-α-damascone/ 4-hydroxy-β-damascone/ 3-oxo-α-ionol | 208 | 370 | 371 | 393 | 369 | 405 | 415 | 739 |
| 3-Hydroxy-α-ionol/ 4-hydroxy-β-ionol/ 3-oxo-7,8-dihydro-α-ionol | 210 | 372 | 373 | 395 | 371 | 407 | 417 | 743 |
| 3-Hydroxy-7,8-dihydro-β-ionol | 212 | 374 | 375 | 397 | 373 | 409 | 419 | 747 |

**Table S2.** UGTs amino acids sequence identities (%).

| | UGT709C6 | UGT86C10 | UGT72AY1 | UGT85A73 | UGT73A25 | UGT73A24 |
|---|---|---|---|---|---|---|
| **UGT709C6** | 100 | 25.3 | 23.6 | 36.1 | 25.9 | 25.5 |
| **UGT86C10** | 25.3 | 100 | 22.1 | 31.1 | 23.9 | 23.9 |
| **UGT72AY1** | 23.6 | 22.1 | 100 | 26.9 | 26.3 | 26.5 |
| **UGT85A73** | 36.1 | 31.1 | 26.9 | 100 | 26.9 | 27.3 |
| **UGT73A25** | 25.9 | 23.9 | 26.3 | 26.9 | 100 | 94.5 |
| **UGT73A24** | 25.5 | 23.9 | 26.5 | 27.3 | 94.5 | 100 |

**Table S3.** The optimal reaction conditions for each UGT for the kinetic assay determined with UDP Glo™ assay.

| | UGT72AY1 | UGT73A25 | UGT73A24 | UGT709C6 | UGT86C10 |
|---|---|---|---|---|---|
| **Substrate** | α-Ionol *1 | α-Ionol | Scopoletin | Carvacrol *2 | Geraniol *3 |
| **Amount (µg)** | 1 | 1 | 0.5 | 0.5 | 1 |
| **Incubation time (min)** | 40 | 10 | 10 | 10 | 10 |
| **Temperature (°C)** | 40 | 40 | 40 | 45 | 35 |
| **pH** | 7.5 | 8.5 | 7.5 | 9.5 | 7 |

*1  Perillyl alcohol and scopoletin were used for the determination of the optimal temperature and pH, respectively

*2  1-Decanol was used for the determination of the optimal temperature and pH

*3  1-Dodecanol was used for the determination of the optimal temperature and pH

**Table S4.** Primers used for amplification of the UGT genes and overexpression qRT-PCR. fwd = forward, rev = reverse.

| Primers | Sequence (5’-3’) | Direction | Purpose | Name/Restriction site |
|---|---|---|---|---|
| *UGT709C6* | CGCGGATCCATGAGGTCTGAAGAAGGAAAAG | fwd | PCR | BamHI |
| | ATAGTTTAGCGGCCGCTCAACCTACCAATGACTTAATATAC | rev | PCR | NotI |
| *UGT86C10* | CGCGGATCCATGGGAGAAATAGAGAAAAATC | fwd | PCR | BamHI |
| | ATAGTTTAGCGGCCGCTCATACTTTTGTTGCACGAAC | rev | PCR | NotI |
| *UGT72AY1* | GAAGATCTATGGATAGCTCACAACTT | fwd | PCR | BglII |
| | CCCTCGAGTTACAACTCTCTGCTCCG | rev | PCR | XhoI |
| *UGT85A73* | CGGGATCCATGGGTTCCATTGGTGCT | fwd | PCR | BamHI |
| | CCCTCGAGTTAATGTTTGGACGAAAG | rev | PCR | XhoI |
| *UGT73A25* | CGGGATCCATGGGTCAGCTCCATATT | fwd | PCR | BamHI |
| | CCCTCGAGTTAATGTCCAGTGGAACT | rev | PCR | XhoI |
| *UGT73A24* | CGGGATCCATGGGTCAGCTCCATTTT | fwd | PCR | BamHI |
| | ATTTGCGGCCGCTTAATGATCAGTAGAACT | rev | PCR | NotI |
| *UGT709C6* | CCTTCATCATCTCCGATTTCAGCCACC | fwd | qRT-PCR | |
| | GAAAGTCATCAGGCCATCAGCGACGT | rev | qRT-PCR | |
| *UGT86C10* | CAGCTCGAGATTCGGGACTCGACATA | fwd | qRT-PCR | |
| | GAAGGTCCCAATGGTACCCCAAGGTAT | rev | qRT-PCR | |
| *UGT72AY1* | CGTGAAGCCTTGCCCAAAAT | fwd | qRT-PCR | |
| | GGATCCACCACGTCATCAGG | rev | qRT-PCR | |
| *UGT73A25* | GCAAGAACCACTGGAACAGC | fwd | qRT-PCR | |
| | CAAACGGAGACACCTGGGTT | rev | qRT-PCR | |
| *UGT73A24* | TGCCGCCCTAATTGTCTTGT | fwd | qRT-PCR | |
| | CTGTCTCTTCCCCAGATCGC | rev | qRT-PCR | |
| *Actin* | CTACGAAGGCTACGCACTCC | fwd | qRT-PCR | |
| | GCAATGTAGGCCAGCTTCTC | rev | qRT-PCR | |
| *IS* | ACCGTTGATTCGCACAATTGGTCATCG | fwd | qRT-PCR | |
| | TACTGCGGGTCGGCAATCGGACG | rev | qRT-PCR | |
